# Nuc domain electrostatics drive the *trans* cleavage activity of CRISPR-Cas12a

**DOI:** 10.1101/2025.10.24.684486

**Authors:** Anthony Newman, Lora Starrs, Gaetan Burgio

## Abstract

The *trans* cleavage activity of type V CRISPR (Clustered Regularly Interspaced Short Palindromic Repeats) Cas12a system has been widely used for the detection of biomolecules. Different Cas12a orthologues exhibit faster or slower *trans* cleavage kinetics, making some orthologues more suited for sensitive molecular detection. Ionic strength of reaction buffers, and mutations that change the electrostatic environment near the RuvC active site have also been reported to strongly influence *trans* cleavage kinetics. Studying three commonly used Cas12a orthologues (FnCas12a, AsCas12a, and LbCas12a), we report that electrostatic interactions near the RuvC active site are critical for their *trans* cleavage activity. Alanine substitution of arginine and lysine residues in the Nuc domain can abolish *trans* cleavage while modestly reducing *cis* cleavage. Substitutions of the RuvC lid and substitutions to introduce positively charged residues in the Nuc could enhance both *cis* and *trans* cleavage. These Cas12a variants improved DNA detection and genome editing efficiency. Overall this study provides a blueprint for future rational engineering of Cas12a nucleases for their *trans* cleavage activities.

**Graphical abstract:** 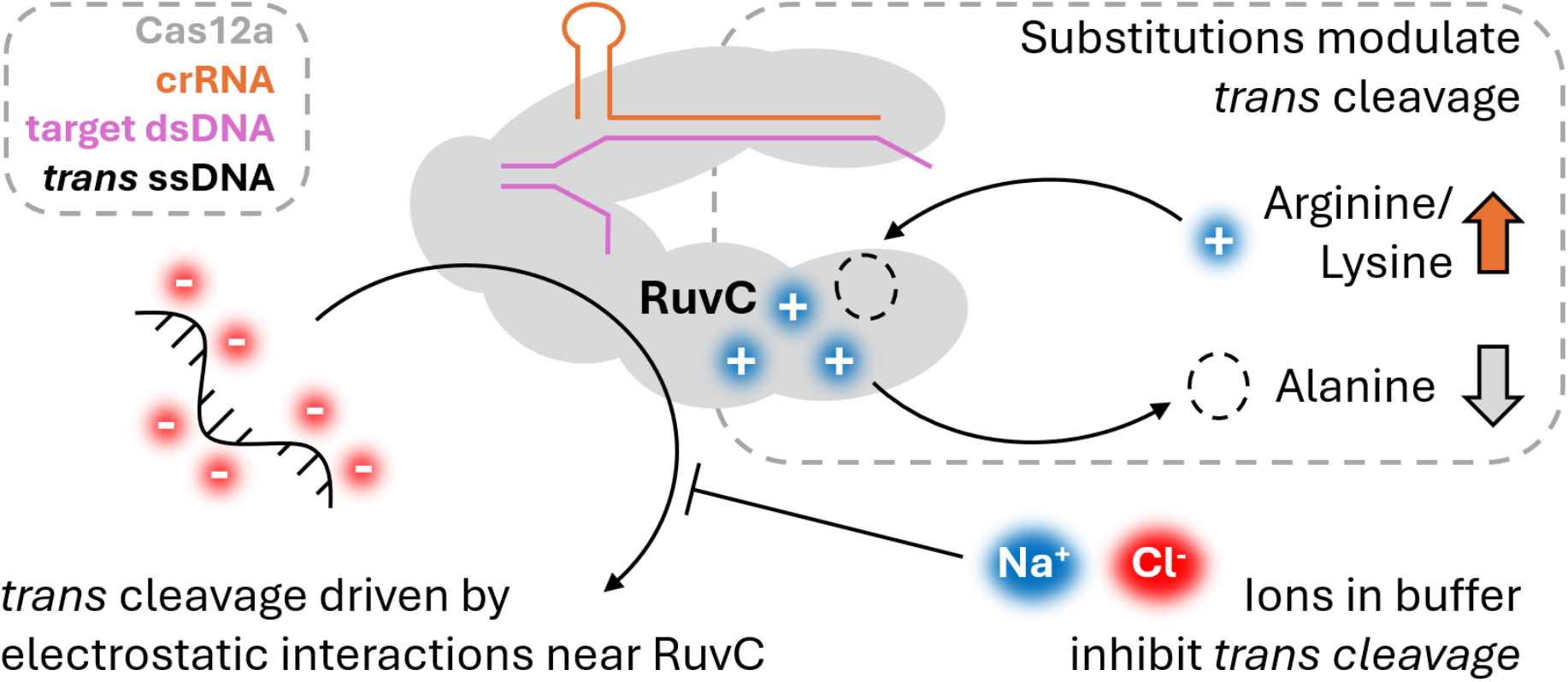

## Introduction

The DNAse activities of the type V-A CRISPR (Clustered Regularly Interspaced Short Palindromic Repeats) Cas12a system have been widely employed for sensitive molecular detection (1). Cas12a has specific RNA guided cleavage of dsDNA (*cis* cleavage), and remains catalytically active to non-specifically cleave ssDNA, RNA, and even nick dsDNA (*trans* cleavage) (2–5). Thus, *trans* cleavage of a suitable reporter molecule signals the presence of the programmed DNA target. Reverse-transcription of RNA can generate a suitable target for *cis* cleavage, and various aptamer strategies can then activate switch *cis* cleavage the presence of proteins, small molecules, and numerous other analytes (6). *Trans* cleavage may then be detected by fluorescence, colorimetry, lateral flow strips, or electrochemical methods (1).

Cas12a binds dsDNA and initiates duplex unwinding at specific nucleotide motifs (the proto-spacer adjacent motif, or PAM) (5). This allows the crRNA guide (CRISPR-RNA) of Cas12a to base-pair to an unwound strand of DNA, the target-strand (or TS) (7, 8). Dynamic motions of the recognition (REC) lobe are stabilised by full matching between the crRNA and TS, which in turn allosterically activates the RuvC active site by stabilising the open state of the RuvC ‘lid’ loop (9, 10). Then, single unwound DNA stands can be coordinated in the active site cleft for cleavage (9, 11). The non-hybridised or non-target DNA strand (NTS) is coordinated by a positively charged groove in the nuclease (NUC) lobe, close to the active site cleft, and in the required 5’ to 3’ polarity for Mg^2+^ mediated phosphodiester bond cleavage (7–9, 11). The scissile phosphate of the TS must twist sharply and traverse **∼**20 Å between the REC and NUC lobes to access the active site, thus rates of TS cleavage are 2-20x slower than NTS cleavage (7–9, 11–16).

Biochemical and structural evidence points to a number of key residues involved in coordinating DNA strands in the active site cleft for cleavage. A conserved arginine in the Nuc domain is adjacent to the RuvC active site residues, and is thought to coordinate the scissile phosphate immediately prior to ‘passing it on’ to the Mg^2+^ ions (7–9, 11, 17). A conserved phenylalanine in the RuvC lid makes base-stacking interactions with incoming substrates, effectively ‘pinning’ DNA strands in the active site (9, 18). Alanine substitution of either residue is deleterious to *cis* and *trans* cleavage (7–9, 11, 17, 18). More broadly, networks of interactions in the Nuc and Nuc-loop also aid coordination of DNA strands for *cis* cleavage (9, 11, 12).

Several studies have identified that *trans* cleavage rates are sensitive to the ionic strength of the reaction buffer (3, 19, 20). For example, increasing NaCl concentration slows *trans* cleavage, and buffers lacking NaCl have higher *trans* cleavage rates (3, 19, 20). Moreover, Cas12a orthologues themselves have varying rates of *trans* cleavage (2, 3, 12). We hypothesised that key electrostatic interactions could be driving *trans* cleavage, and that these vary between Cas12a orthologues.

To test this hypothesis, we performed alanine substitution of positively charged residues in the Nuc across three Cas12a orthologues (FnCas12a from *Francisella tularensis subsp. novicida U112*, AsCas12a from *Acidaminococcus sp. BV3L6*, and LbCas12a from *Lachnospiraceae bacterium ND2006*). We found that alanine substitution could abolish *trans* cleavage, whilst retaining *cis* cleavage. This complements literature observations of ionic strength in reaction buffers decreasing *trans* cleavage rates, by neutralising non-specific electrostatic protein-nucleic acid interactions (19). To engineer Cas12a nucleases with higher *cis* and *trans* cleavage, we assess mutations in the RuvC-lid, and perform substitutions to introduce more positively charged residues in the Nuc domain. These variants exhibited higher *trans* cleavage, especially at higher NaCl concentrations, enhancing their DNA detection ability. Engineered variants also have increased *cis* cleavage kinetics, and improved genome editing activity.

## Methods

### Structural analyses

Structural data of Cas12a orthologues was accessed from PDB (rcsb.org), or EMBL/AlphaFold databases (6GTG and 6I1K – FnCas12a, 8SFR – AsCas12a, A0A182DWE3 – LbCas12a) (21). Structures were visualised in ChimeraX (version 1.9) (22).

### Cloning, protein purification, and protein thermostability

Site-directed mutagenesis was used to generate point mutants of Cas12a orthologues. Template plasmids were pET21a-Cas12a-3xHA-2xNLS constructs, as used previously (12). Q5 site-directed mutagenesis was performed as per the manufacturer’s instructions (NEB), using primers in **Table S1**. Sequences were verified by Sanger sequencing (performed at the Biomolecular Resource Facility, ANU). Cas12a proteins were obtained by overexpression and purification from T7Express cells (NEB), as previously described, with no modifications (12). Likewise, protein thermostability assays were performed precisely as previously described (12).

### Trans cleavage kinetics

Oligonucleotides for crRNA and DNA targets (**Table S2**) were purchased from Integrated DNA Technologies (IDT), and resuspended in 1x IDTE buffer (IDT). Double-stranded DNA targets were made by annealing 10 µM each TS and NTS oligos in 1x Duplex Buffer (IDT), and heating to 90 °C in a thermocycler, and cooling at 1 °C per 30s until 20 °C. DNA targets for *cis* cleavage (1 nM, unless otherwise indicated) were prepared in cleavage buffer (10 mM Tris-HCl, pH 7.5, 10 mM MgCl_2_, 5 µg/ml bovine serum albumin [BSA], 50 mM NaCl - unless otherwise indicated), with 1 mM of freshly prepared dithiothreitol (DTT).

Cas12a protein and crRNA were complexed by directly mixing crRNA (in IDTE buffer) with Cas12a (in storage buffer, 20 mM Tris-HCl, pH 7.5, 500 mM NaCl, 50% glycerol, 1 mM DTT), and incubated at room temperature for 10 mins. *Cis* cleavage was performed by addition of DNA target solution, to a final concentration of 10 nM Cas12a-crRNA and 1 nM target dsDNA, in 1x cleavage buffer (NaCl concentration as indicated). This *cis* cleavage reaction was incubated in a thermocycler at 30 °C for strictly 20 mins.

This ternary Cas12a-crRNA-target DNA complex was then further diluted in appropriate 1x cleavage buffer, and 50 uL added to each well of a flat-clear-bottom black fluorescence 96-well plate (ThermoFisher). Fluorescent-quencher reporter ssDNA was prepared in 1x cleavage buffer (NaCl as indicated, reporter ssDNA final concentration 75 nM), 50 µL of which was added into each well by multichannel pipette. *Trans* cleavage in the absence of target DNA or crRNA was performed in no-NaCl cleavage buffer, and 500 nM reporter ssDNA. Fluorescence over time for all assays was measured in a Victor Nivo (PerkinElmer) or an Infinite M Nano plate reader (Tecan), using the 480/30nm filter for excitation and 530/30nm filter for emission.

Reactions with a final concentration of ∼1 mM NaCl were attained by a total of 500-fold dilution across the steps of crRNA complexing, *cis* cleavage reaction, and addition of reporter ssDNA; the Cas12a proteins being stored in a buffer containing 500 mM NaCl.

Fluorescence curves were plotted GraphPad Prism 10 (GraphPad Software). For pseudo-first order *trans* cleavage kinetics, the first 300s were extracted and a linear regression fitted to each replicate (GraphPad Software). Values of fluorescence increase per second (ΔFs^-1^) were then plotted (GraphPad Software).

### Cis cleavage kinetics

These were performed as previously described (12), with the exception of FnCas12a WT kinetics performed in cleavage buffers containing final concentration of ∼1, 100, or 200 mM NaCl, as indicated.

Briefly, Cas12a-crRNA complexes (cognate DNMT1-3 crRNA, **Table S2**) were assembled and incubated with a target plasmid for set time intervals. The obligatory sequential NTS then TS cleavage mechanism of Cas12a causes changes in plasmid DNA topology, from negatively supercoiled, to the ‘nicked’ open-circle, and finally to the linearised form (12, 14, 15). These changes in topology are visible by agarose gel electrophoresis (23), and supercoiled, nicked, and linearised fractions were quantified by ImageJ (24). Fitting of a sequential strand cleavage model to changes in DNA topology over time was used to derive rate constants of NTS and TS cleavage (12, 14, 15).

### DNA detection experiments

Cas12a-crRNA complexes were prepared as for *trans* cleavage kinetics. Complexes were diluted in 1x cleavage buffer to 10 nM (final NaCl concentration as indicated), and 10 µL added per well (Applied Biosystems MicroAmp Optical 96-well Reaction Plate, ThermoFisher). ‘DNA-reporter’ solution was prepared, containing a final concentration of 200 mM fluorescent-quencher reporter ssDNA and 50 pM target dsDNA, in 1x cleavage buffer (final NaCl concentration as indicated). 10 µL DNA solution was added to the 96-well plate (on ice) using a multichannel pipette and mixed well. The plate was covered with adhesive film (MicroAmp Optical, ThermoFisher), and loaded into an Applied Biosystem quantitative-PCR machine. Fluorescence values were recorded every 30 s, and curves plotted in GraphPad Prism 10 (GraphPad Software). Statistical tests on endpoint fluorescence values at 30 minutes were performed to assess DNA detection ability, using two-way ANOVA followed by Tukey’s multiple comparisons test in GraphPad Prism 10 (GraphPad Software).

DNA detection in human saliva was performed as above, with the following modifications. The DNAse activity of saliva was inactivated by addition of 2% v/v Proteinase K (Bioline), with incubation at 55 ºC for 10 mins, then 98 ºC for 10 mins. A final concentration of 150 nM ssDNA reporter was used, and the remaining 63% of sample volume consisted of human saliva (Innovative Research Inc Pooled Human Saliva 5ml, ThermoFisher).

### Genome editing of cell lines

Three replicate editing experiments were performed per nuclease, precisely as previously described (12) – with the following addition. Positive control consisting of 30 pM Alt-R AsCas12a Ultra (IDT) was complexed with 0.575 µM DNMT1-3 crRNA, and complexed as previously described (12). This was then delivered by electroporation into HEK293T, A549, and Jurkat cells. Cell culturing, electroporation, genomic DNA extraction, and high-throughput sequencing were then performed precisely as described (12). Statistical significance of gene editing was performed on percentage of indels, using two-way ANOVA followed by Tukey’s multiple comparisons test in GraphPad Prism 10 (GraphPad Software).

## Results

### Alanine substitution near the RuvC can abolish the *trans* cleavage of Cas12a orthologues

We first generated alanine substitutions of a conserved arginine in the Nuc domain (FnR1218, AsR1226, and LbR1138, **Figure 1 A,B,C**) that is critical for *cis* and *trans* cleavage (4, 7, 8). We determined the *cis* cleavage kinetics of these mutants, and replicated previous results showing globally depressed rates of NTS and TS cleavage, with no detectable *trans* cleavage activity (**Figure S1**) (4, 8, 11, 25).

**Figure 1:**
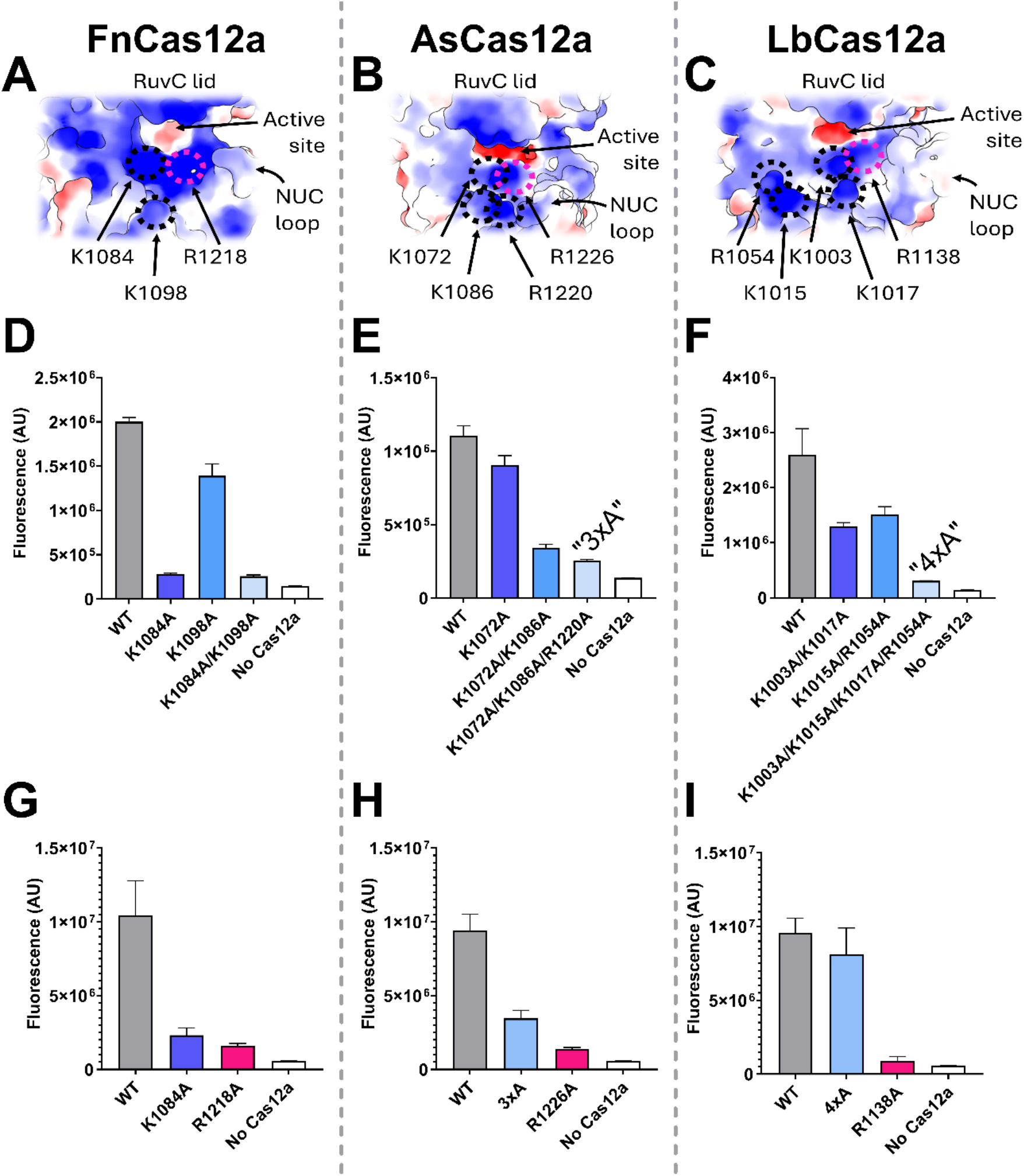
Coulombic electrostatic potential mapped onto the surface of (**A**) FnCas12a [6GTG], (**B**) AsCas12a [8SFR], and (**C**) LbCas12a [A0A182DWE3 -AFDB prediction to show incomplete lid and Nuc-loop]. Electrostatic potential is displayed on a colour gradient showing negative (red), neutral (white), and positive (blue). Arrows show the RuvC active site cleft, Nuc loop, and key amino acids that contribute to positive surface charge. Endpoint fluorescence after 60 mins *trans* cleavage reaction, when activated with 1nM target dsDNA, for wild-type and mutants of (**D**) FnCas12a, (**E**) AsCas12a, and (**F**) LbCas12a. Endpoint fluorescence after 60 mins *trans* cleavage reaction, when activated with 1nM ‘pre-cleaved’ dsDNA substrate, for WT and mutants of (**G**) FnCas12a, (**H**) AsCas12a, and (**I**) LbCas12a.

We next wondered if other nearby positively charged residues had similar effects on DNA cleavage. We identified a lysine residue (FnK1084, AsK1072, LbK1003), adjacent to the conserved arginine (FnR1218, As R1226, LbR1138). Mapping the Coulombic electrostatic potential onto the RuvC and Nuc domains showed this amino acid, and several others, contribute to the positive charge of the protein surface near the active site cleft (**Figure 1A,B,C**, **Figure S2-4**).

Alanine substitution of K1084 in FnCas12a was sufficient to reduce *trans* cleavage to near undetectable (**Figure 1D, Figure S5A**). This was not the case for the orthologous substitution in AsCas12a (K1072A, **Figure 1E, Figure S5B**), nor in LbCas12a (K1003A, **Figure S5C**). We generated single, double, triple, and quadruple alanine substitutions to achieve Cas12a mutants with severely reduced *trans* cleavage activity (**Figure 1D,E,F**). For AsCas12a, triple alanine substitution mutant ‘3xA’ (K1072A/K1086A/R1220A) exhibited the weakest *trans* cleavage activity (**Figure 1E, Figure S5B**), for LbCas12a it was quadruple alanine substitution mutant ‘4xA’ (K1003A/K1015A/K1017A/R1054A, **Figure 1F, Figure S5C,D**). We next assayed the *cis* cleavage kinetics of these mutants (**Figure S6, Table S4**). NTS and TS cleavage of FnCas12a K1084A reduced by 1.8- and 1.3-fold, while AsCas12a ‘3xA’ reduced 2.1- and 7-fold respectively (**Figure S7, Table S4**). NTS cleavage of LbCas12a mutants was most affected, reduced 1.5-to 2.1-fold from double to quadruple alanine substitution mutant (**Figure S7, Table S4**). TS cleavage was less affected, slowed by 1.1-to 1.5-fold (**Figure S7, Table S4**).

We reasoned that these alanine substitution mutants lose *trans* cleavage activity through loss of electrostatic interactions between positive charges on the surface of the Nuc domain and the negatively charged backbone of substrate DNA. This is consistent with numerous reports showing decreased *trans* cleavage rates with increasing ionic strength in reaction buffers (3, 19, 20). We replicated this effect, in reaction buffers ranging from ∼1 to 100 mM NaCl (**Figure S8,9**). Here, the higher NaCl concentration slows *trans* cleavage, and lower NaCl concentrations allow increasing rates of *trans* cleavage.

This effect varied between Cas12a orthologues, with LbCas12a the least affected by increasing NaCl concentration (**Figure S8,9**). This effect is consistent with our mutagenesis study, where LbCas12a required four alanine substitutions to abolish *trans* cleavage, while for FnCas12a and AsCas12a one and two substitutions were sufficient to suppress most *trans* cleavage activity (**Figure 1D,E,F**). Given a single alanine substitution could effectively abolish the *trans* cleavage of FnCas12a, and *trans* cleavage could be well suppressed by high NaCl concentration, we explored the effect of NaCl concentration on the *cis* cleavage kinetics of FnCas12a. We found that *cis* cleavage rates did not greatly change across 1-200 mM NaCl, with *k*_NTS_ and *k*_TS_ varying by 1.6 to 1.3-fold respectively (**Figure S10,11, Table S4**). More important was alanine substitution of residues near the RuvC, with the FnK1084A/K1098A double mutant having 1.7-fold reduced *k*_NTS_ and 2.7-fold slower *k*_TS_ (**Figure S11, Table S4**).

To further test our defectively *trans* cleaving Cas12a mutants, we complexed them with a ‘pre-cleaved’ dsDNA substrate containing a 20 nt TS and 14 nt NTS, and tested their *trans* cleavage in the low salt condition (∼1 mM NaCl, **Figure 1G-I, Figure S12**). This assay showed that single mutants K1084A and R1218A of FnCas12a have very little affinity to the ssDNA *trans* substrate (**Figure 1G,H, Figure S12B,C**). AsCas12a mutant ‘3xA’ retained very weak *trans* cleavage, while the LbCas12a ‘4xA’ retained the most *trans* cleavage activity (**Figure 1H, Figure S12D**). The ‘4xA’ mutant had a significantly slower increase in fluorescence, despite reaching similar values to WT LbCas12a after 60 mins, indicating this mutant was significantly impaired in its *trans* cleavage activity.

That this ‘4xA’ mutant of LbCas12a retained some degree of *trans* cleavage suggested that the LbR1138 residue alone could mediate this activity, unlike the R1218 of FnCas12a or R1226 of AsCas12a. In the allosteric activation of the RuvC active site, the occluding lid loop opens and closes (9). We reasoned that LbCas12a may have intrinsically faster rates of ‘opening’ the RuvC lid, relative to FnCas12a and AsCas12a. The lid contains a conserved phenylalanine that base-stacks with incoming nucleic acids (Fn F1012, As F999, Lb F931), and aids their coordination in the RuvC alongside the network of positively charged Nuc residues. So, we analysed the structures of the RuvC lid of Cas12a orthologues.

### Lid and Nuc substitutions can increase the *trans* cleavage activity of Cas12a orthologues

In analysing structures of Cas12a, we noted that a lid-NTS stacking interaction was present in the closed-lid conformation of FnCas12a (**Figure 2A**). Here, F1010 of FnCas12a forms a stacking interaction with the 11^th^ nucleotide of the non-target DNA strand (11). This stacking between F1010 and the NTS was not present in structures showing the ‘open’ lid conformation (**Figure 2A**) (26). Alignment of the RuvC lid between Cas12a orthologues showed that LbCas12a harbour a serine in this position, unlike the phenylalanine of FnCas12a and AsCas12a. We hypothesised that removing the possibility of lid-NTS stacking may promote the open lid conformation, increasing *trans* cleavage.

**Figure 2:**
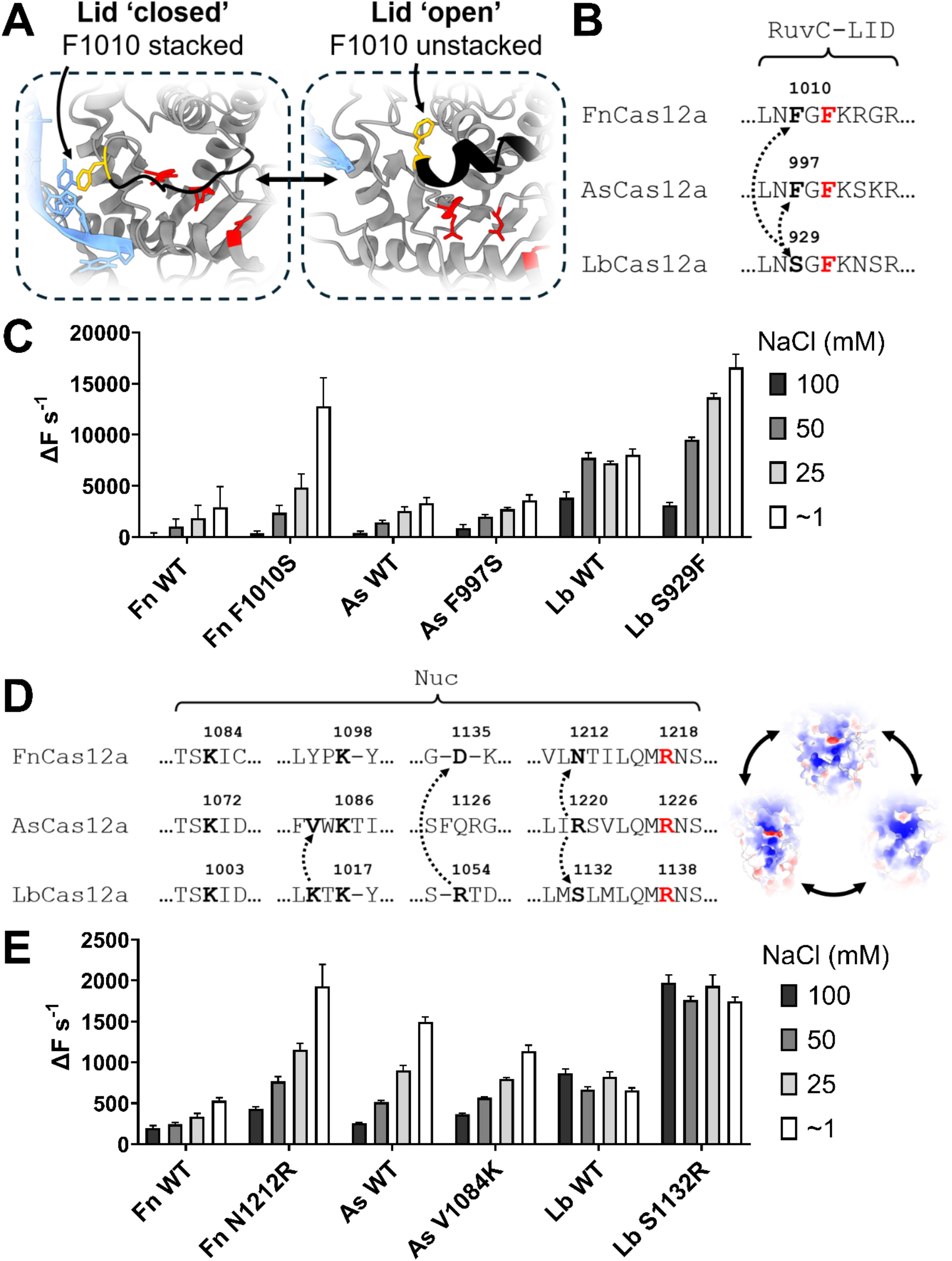
(**A**) Diagram of the RuvC active site of FnCas12a, catalytic residues shown in red. Polypeptide is shown in grey, while Lid motif is shown in black. Left box shows the RuvC lid of FnCas12a in the closed state (PDB: 6I1K), and the right box in the open state (PDB: 6GTG). Residue F1010 (mustard yellow) is shown interacting with the non-target DNA strand (blue). **(B)** Alignment of RuvC lid of FnCas12a, AsCas12a, and LbCas12a, with conserved lid – phenylalanine residue in bold red. Amino acids substituted in bold. **(C)** *Trans* cleavage rates of WT Cas12a and mutants, as quantified by the initial slope of fluorescence curve. *Trans* cleavage reactions performed in buffers with final concentration of NaCl from ∼1 to 100 mM. **(D)** Schematic of orthologue-informed mutagenesis. Selected alignment of Nuc domains of FnCas12a, AsCas12a, and LbCas12a, with mutated residues in bold. Conserved RuvC-proximal arginine in red. **(E)** *Trans* cleavage rates of WT Cas12a and mutants, as quantified by the initial slope of fluorescence curve. *Trans* cleavage reactions performed in buffers with final concentration of NaCl from ∼1 to 100 mM.

To test this hypothesis, we generated orthologue-informed substitutions, whereby the serine of LbCas12a replaced the phenylalanine of FnCas12a and AsCas12a, and vice versa (**Figure 2B**), and tested their *trans* cleavage in the ∼1 mM to 100 mM NaCl conditions. FnCas12a F1010S had increased trans cleavage relative to WT, which was more pronounced at ∼1 and 25 mM NaCl (**Figure 2C, Figure S13**). AsCas12a F997S exhibited similar rates of *trans* cleavage to WT (**Figure 2C, Figure S14**). We expected the LbCas12a mutant S929F would show decreased cleavage activity, however, we observed increased *trans* cleavage rates (**Figure 2C, Figure S15**). The increase in *trans* cleavage was notable in the ∼1 and 25 mM NaCl condition, and not apparent at 50 and 100 mM NaCl (**Figure 2C, Figure S15**).

Encouraged by the possibility of engineering Cas12a nucleases with increased *trans* cleavage, we generated orthologue-informed substitutions of the Nuc domain, this time to introduce positively charged amino acids where an orthologue was lacking (**Figure 2D**). We chose positions shown to be important for *trans* cleavage in **Figure 1**. For example, residue R1220 of AsCas12a aligns structurally with FnCas12a N1212 and LbCas12a S1132. Likewise, LbCas12a R1054 aligns with AsCas12a V1084. Thus, we substituted these amino acids for arginine or lysine.

We generated mutations of the Nuc domain, and tested their *trans* cleavage across the ∼1 mM to 100 mM NaCl conditions. Mutation N1212R for FnCas12a improved *trans* cleavage rates across NaCl concentrations, while V1084K substitution for AsCas12a showed a very slight improvement at 100 mM NaCl (**Figure 2E, Figure S16,17**). For LbCas12a, the S1132R mutation consistently improved *trans* cleavage rates across NaCl concentrations (**Figure 2E, Figure S18**). Substitution mutant D1135K for FnCas12a exhibited *trans* cleavage rates comparable to WT in ∼1 to 50 mM, and slightly increased at 100 mM NaCl (**Figure S19**). Double mutants FnF1010S/N1212R and FnD1135K/N1212R showed increased *trans* cleavage, while the triple mutant F1010S/D1135K/N1212R, or ‘SKR’, showed the highest *trans* cleavage activity (**Figure S20**).

### Engineered Cas12a nucleases show enhanced *cis* cleavage and DNA detection

We next assayed the *cis* cleavage kinetics of single and combination mutants (**Figure S21,S22**). Single mutants of FnCas12a rates of NTS and TS cleavage similar to WT, while double mutants F1010S/N1212R and D1135K/N1212R had decreased rates of *k*_TS_ and *k*_NTS_ respectively (**Figure 3A, Table S4**). Triple mutant ‘SKR’ showed increased *k*_NTS_ relative to WT, and similar *k*_TS_ (**Figure 3A, Table S4**). AsCas12a F997S had similar *cis* cleavage kinetics to WT, while the V1084K mutation decreased *k*_NTS_ by 4.5-fold, and increased *k*_TS_ by 1.6-fold (**Figure 3B, Table S4**). LbCas12a S929F mutant exhibited 2.5-fold increased *k*_NTS_ relative to WT, while S1132R substitution increased *k*_TS_ by 2.6-fold (**Figure 3B, Table S4**). Combination mutant S929F/S1132R increased *k*_NTS_ by 2.1-fold and *k*_TS_ by 1.3-fold (**Figure 3B, Table S4**). We further assessed the effect of mutations on protein thermostability, which determined single and combination mutants of Cas12a orthologues have similar-to-WT stability (**Figure S23**).

**Figure 3:**
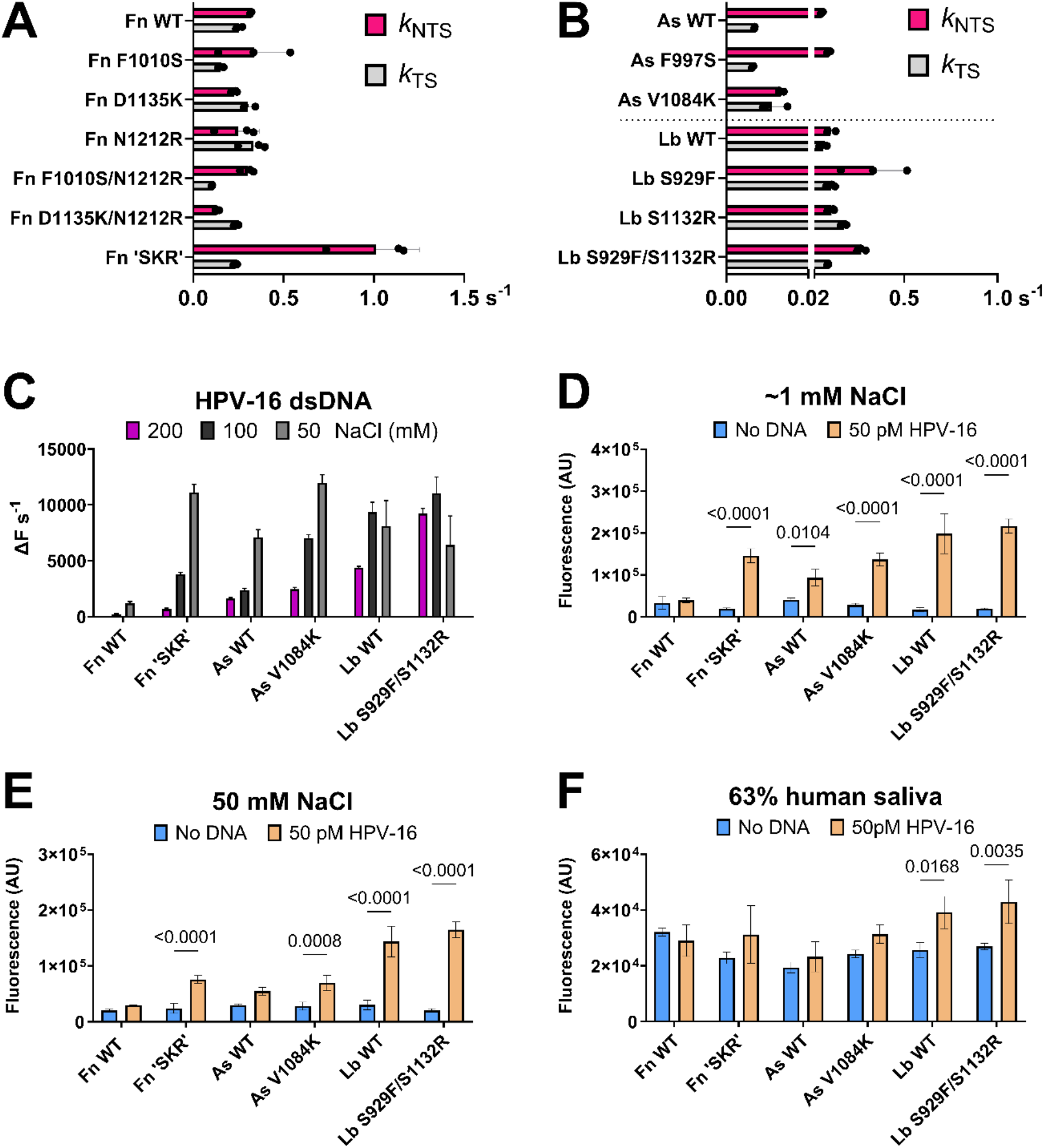
(**A, B**) Kinetic values (s^-1^) of NTS (*k*_NTS_, pink) and TS cleavage (*k*_TS_, grey), for the Cas12a indicated. Dots show individual replicates, bars show mean, error bars show s.d. **(C)** *Trans* cleavage rates with HPV-16 dsDNA target, in 50, 100, and 200 mM NaCl, as quantified by the initial slope of fluorescence curve. **(D)** Detection of 50 pM HPV-16 dsDNA, by endpoint fluorescence at 30 minutes, in ∼1 mM final concentration of NaCl. **(E)** Detection of 50 pM HPV-16 dsDNA, by endpoint fluorescence at 30 minutes, in 50 mM final concentration of NaCl. **(F)** Detection of 50 pM HPV-16 dsDNA, by endpoint fluorescence at 30 minutes, in sample that is 63% (v/v) human saliva. Statistical significance evaluated by two-way ANOVA with Tukey’s multiple comparison test, p-values shown above bars.

Engineered Cas12a mutants have additional positive charges in their Nuc domains. We hypothesised these mutants would show increased *trans* cleavage at higher ionic strengths. We tested this hypothesis using an HPV-16 crRNA and matching dsDNA target, across 50-200 mM NaCl (**Figure 3C, Figure S24**). FnCas12a triple mutant ‘SKR’ showed increased *trans* cleavage rates at 50 to 100 mM NaCl, with both WT and mutant have very slow *trans* cleavage at 200 mM (**Figure 3C, Figure S24**). AsV1084K had similar *trans* cleavage rates to WT at 200 mM NaCl, and increased at 50 to 100 mM (**Figure 3C, Figure S24**). Double mutant LbS929F/S1132R had similar to WT rates of *trans* cleavage, but increased at the 200 mM condition (**Figure 3C, Figure S24**).

We next tested these engineered Cas12a nucleases for their ability to detect low concentrations of DNA. To illustrate the effect of monovalent ions on *trans* cleavage, we conducted this assay in both ∼1 mM and 50 mM NaCl conditions, with the low-salt condition generally resulting in higher endpoint fluorescence (**Figure 3D-E, Figure S25,26**). Engineered nucleases typically out-performed wild-types, with the exception of LbCas12a WT and S929F/S1132R, which both displayed very robust DNA detection (**Figure 3D-E, Figure S25,26**). Unlike FnCas12a and LbCas12a nucleases, AsCas12a WT and mutants had relatively high rates of *trans* cleavage in the absence of crRNA or target DNA, limiting their DNA detection sensitivity (**Figure S27**). Consistent with previous reports, we found this activity is mediated by *apo* AsCas12a, as it decreases with molar excesses of crRNA (**Figure S28**). Finally, we performed DNA detection in a sample matrix of 63% human saliva (**Figure 3F, Figure S29,30**). Endpoint fluorescence values were lower, and only LbCas12a WT and S929F/S1132R had significantly increased fluorescence compared to the no-DNA control, with S929F/S1132R out-performing WT LbCas12a (**Figure 3F, Figure S30**).

Finally, we tested the genome editing efficiency of high- and low-*trans* cleaving Cas12a mutants of AsCas12a and LbCas12a. We chose three commonly used human cell lines – HEK293T, A549, and Jurkat. We electroporated Cas12a complexed to crRNA targeting either the DNMT1-3, DNMT1-7, or AGBL1 genes, and insertions or deletions (indels) were assessed by high-throughput sequencing.

Cas12a mutants with low-*trans* cleavage typically showed reduced indels, while high-*trans* cleavage mutant had retained or increased editing relative to WT (**Figure 4**). AsCas12a triple mutant ‘3xA’ had significantly reduced indel activity at the DNMT1-3 site, across all three cell lines tested. Mean indels for WT and mutants were lower overall at the DNMT1-7 and AGBL1 sites, but the ‘3xA’ mutant had consistently lower-than-WT editing activity (**Figure 4A, C, E**). AsV1084K had editing efficiencies similar to WT across all cell lines and target sites, and significantly higher at the DNMT1-3 site in HEK293T cells (**Figure 4A**). The LbCas12a ‘4xA’ mutant had similar or lower editing efficiency to WT LbCas12a, while S929F/S1132R mutant had similar or increased activity (**Figure 4B, D, F**). This increase was only weakly significant at the DNMT1-7 site in Jurkat cells (**Figure 4F**). We also assessed editing at previously identified off-target sites (**Figure S31-33**) (27). All nucleases showed no significant increase in off-target editing – with the exception of AsV1084K – which exhibited increased editing at off-target site 1 with the DNMT1-3 crRNA, in A549 and Jurkat cells (**Figure S32,33**). This off-target editing reached a maximum of 3% indels (**Figure S32**).

**Figure 4:**
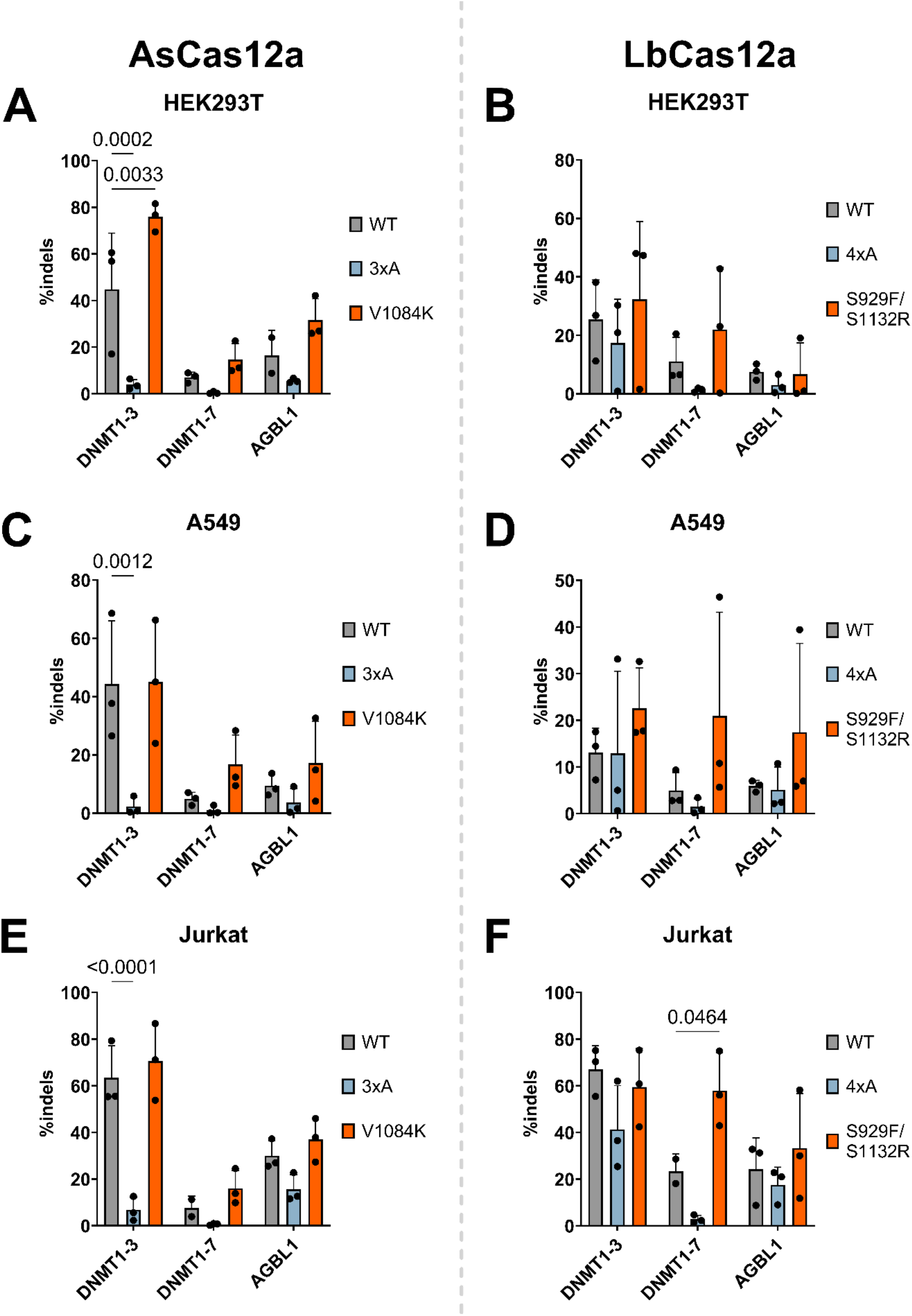
Genome editing in HEK293T cells by (**A**) AsCas12a and (**B**) LbCas12a nucleases. Genome editing in A549 cells by (**C**) AsCas12a and (**D**) LbCas12a nucleases. Genome editing in Jurkat cells by (**E**) AsCas12a and (**F**) LbCas12a nucleases. Statistical significance evaluated by two-way ANOVA with Tukey’s multiple comparison test, p-values are shown above bars.

We further compared our engineered nucleases by comparison with a commercially available Cas12a variant – AsCas12a ‘Ultra’ (28). We compared on- and off-target editing using the DNMT1-3 crRNA across three cell lines (**Figure S34,35**). AsCas12a ‘Ultra’ had significantly higher on-target indel activity than our nucleases in HEK293T and A549 cell lines, with similar but lower activity in Jurkat cells (**Figure S34**). However, AsCas12a ‘Ultra’ exhibited very high activity at the DNMT1-3 off-target-1, with 20 to 70% indels across all three cell lines tested (**Figure S35**).

## Discussion

Cas12a *trans* cleavage activity has been widely employed for molecular detection. However, its mechanism and kinetics remains elusive. We hypothesise electrostatic interactions plays a critical role in *trans* cleavage activity of Cas12a orthologues. We explored the role of Nuc and RuvC lid mutation on Cas12a *trans-cleavage* activity. Our study shows that electrostatic interactions are key drivers of the *trans* cleavage activity of Cas12a orthologues. This finding rationalises the high *trans* cleavage activity seen with low-salt buffers, and allows engineering of improved Cas12a variants for molecular detection.

### Nuc electrostatics are critical for the *trans* cleavage of Cas12a orthologues

We replicated previous findings that FnR1218A, AsR1226A, and LbR1138A had highly impaired *cis* cleavage kinetics (3 to 6 orders of magnitude slower than WT), and no detectable *trans* cleavage (**Figure S1**). We further determined that these mutants were unable to effect *trans* cleavage even when complexed to a pre-cleaved dsDNA substrate, in a no-NaCl reaction buffer (**Figure 1G,H,I**). This indicates this residue provides critical affinity to *trans* substrate ssDNA (**Figure 5**).

**Figure 5:**
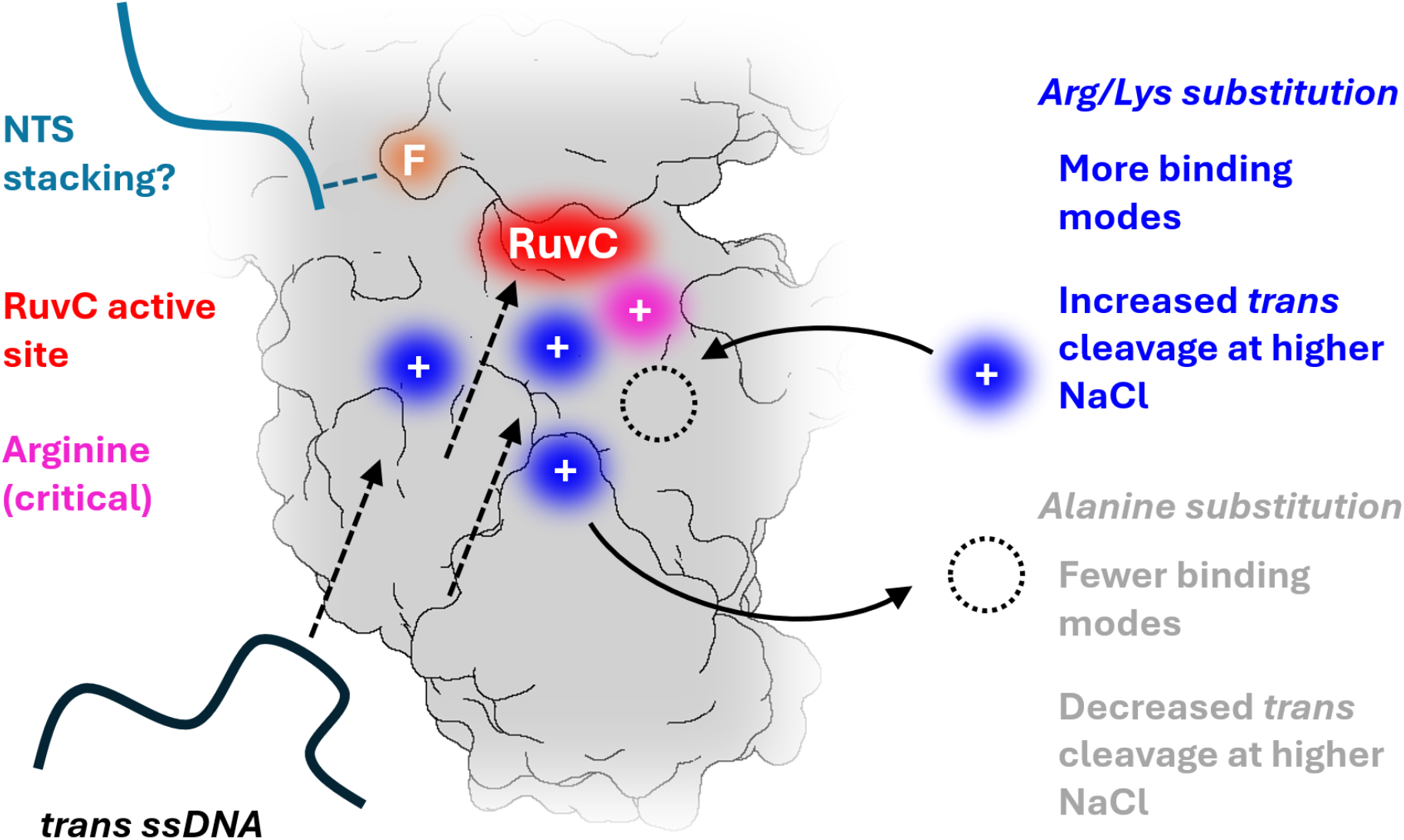
Model of Nuc electrostatics and *trans* cleavage, showing; putative stacking interaction between NTS (sea green) and RuvC lid phenylalanine (orange), RuvC active site cleft (red), with adjacent conserved and critical arginine residue (pink), positively charged residues (blue) that play a role in *trans* cleavage, which can be removed by alanine substitution (dashed circle), and *trans* ssDNA substrate (black) with arrows showing potential modes for binding and cleavage by Cas12a. NUC lobe of FnCas12a (grey, PDB: 6GTG) is used as a generic representation, as are the positions of residues and nucleic acid substrates.

Alanine substitution of other lysine and arginine residues near the RuvC had variable effects on *cis* and *trans* cleavage across Cas12a orthologues (**Figure 1D,E,F**, **Figure S5-7**). This is likely due to different exact protein-nucleic acid interactions Cas12a orthologues employ to bind and cleave *cis* and *trans* substrates. The NTS and TS are coordinated to Cas12a by numerous interactions across the REC and NUC lobes (7–9, 11, 17, 26), while *trans* substrate ssDNA is predicted to transiently interact with Nuc and RuvC domains (29). We propose that this weaker interaction between *trans* substrate ssDNA and Cas12a underpins the greater loss of *trans* cleavage vs *cis* cleavage upon alanine substitution of Nuc residues -i.e. the single mutant FnK1084A abolishes *trans* cleavage while retaining *cis* cleavage rates within 2-fold of WT (**Figure S5A, FigureS7**). Similarly, changing NaCl concentrations in reaction buffers from 1-200 mM NaCl had relatively little effect on the *cis* cleavage of FnCas12a WT, unlike observed for *trans* cleavage (**Figure S8-11**).

AsCas12a single mutants K1072A and K1086A reduced *trans* cleavage rates relative to WT, the double mutant further reducing *trans* cleavage, and triple combination mutant with R1220A – ‘3xA’ – having the weakest *trans* cleavage activity (**Figure S5B**). We suggest that R1226, K1072, and K1086 are the major contributors to *trans* substrate binding, with some contribution of R1220. We suggest these residues also contribute significantly to *cis* substrate binding, the ‘3xA’ mutant having 2.1-fold lower *k*_NTS_ and 7-fold slower *k*_TS_ compared to AsCas12a WT (**Figure S7**).

LbCas12a single mutants of K1003A and R1054A reduced *trans* cleavage to a greater extent than K1015A and K1017A (**Figure S5C**). Double mutants including K1003A were more impaired in trans cleavage than double mutants not including it – i.e. K1015A/K1017A and K1015A/R1054A (shown in **bold, Figure S5C**). This would suggest *trans* ssDNA substrates bind to R1138 and K1003 in conjunction with other positively charged residues – i.e. K1015, K1017, or R1054. The LbCas12a ‘4xA’ mutant retained a degree of *trans* cleavage when incubated with a pre-cleaved *cis* substrate, compared to the minimal activity of FnCas12a K1084A and AsCas12a ‘3xA’ (**Figure S12**). This would suggest the LbCas12a R1138 residue alone has reasonable affinity to *trans* substrate ssDNA. Indeed, despite the slower *cis* cleavage kinetics of LbCas12a ‘4xA’ (**Figure S7**), this mutant displayed similar-to-WT levels of gene editing at some target sites, unlike the AsCas12a ‘3xA’ mutant (**Figure 4**).

Our alanine mutagenesis study of FnCas12a, AsCas12a, and LbCas12a indicates that electrostatic interactions of Nuc domain residues are critical for the *trans* cleavage of these orthologues. Specifically, alanine substitution of positively charged residues closest to the active site cleft tend to have a larger negative effect on DNAse activity, as does alanine substitution of multiple such residues of the Nuc.

### RuvC-lid and Nuc substitutions can generate Cas12a variants with enhanced *cis* and *trans* cleavage

We explored the role of RuvC-lid substitution in the *cis* and *trans* cleavage of Cas12a orthologues. It had been previously established that a conserved phenylalanine of the RuvC-lid makes key interactions with *cis* and *trans* substrates (FnF1012, AsF999, LbF931) (9, 18). Alanine substitution at this position reduced the TS cleavage kinetics of FnCas12a and AsCas12a (18), and abolished *trans* cleavage for AsCas12a (9). We tested a less-conserved lid residue seen to interact with the NTS (FnF1010, AsF997, LbS929) (11), and found inconsistent effects on *cis* and *trans* cleavage (**Figure 2C**). Substitution mutant LbS929F exhibited increased *cis* and *trans* cleavage, as did FnF1010S to a lesser extent, while AsF997S had little effect on *cis* and *trans* cleavage (**Figure 3A,B**). These data make the mechanistic role of this position unclear.

Nuc substitution had unpredictable effects on the *cis* cleavage of FnCas12a. While single substitution mutants of FnCas12a had similar *cis* cleavage kinetics to WT, double mutant FnD1135K/N1212R had 2.4-fold decreased *k*_NTS_ and FnF1010S/N1212R had 2.5-fold decreased *k*_TS_ (**Figure 3A**). However, triple mutant FnF1010S/D1135K/N1212R – ‘SKR’ – exhibited ∼3-fold increased *k*_NTS_ and enhanced *trans* cleavage kinetics (**Figure 3A**).

AsCas12a V1084K mutant had 1.6-fold increased *k*_TS_ with 4.6-fold lower *k*_NTS_ (**Figure 3B**). This mutation may interfere with NTS-loading into the RuvC, by stabilising DNA binding further from the active site. AsV1084K slightly increased *trans* cleavage in higher NaCl conditions, having a greater effect when activated by the HPV-16 dsDNA target (**Figure 2E, 3C**). The LbS1132R mutant increased *k*_TS_ by 2.7-fold, with similar *k*_NTS_ (**Figure 3B**), and consistently increased *trans* cleavage rates across NaCl conditions (**Figure 2E**). The combination mutant LbS929F/S1132R had 2.6-fold increased *k*_NTS_, 1.4-fold increased *k*_TS_ (**Figure 3B**), with enhanced *trans* cleavage in higher NaCl concentrations (**Figure 3C**).

In total, these data suggest that orthologue-informed substitutions can be used to enhance the DNAse activities of Cas12a. Previous structure-guided engineering of Cas12a has focussed on introducing positively charged amino acids to enhance PAM or crRNA-TS heteroduplex interactions (27, 30). Here, we show that RuvC-lid and Nuc substitutions can enhance both *cis* and *trans* cleavage activity FnCas12a, AsCas12a, and LbCas12a, suggesting this approach could be applicable to other Cas12a orthologues.

### High-trans cleaving Cas12a variants can enhance DNA detection and genome editing

We tested high-*trans* cleaving variants of Cas12a for their ability to detect low amounts of target DNA. In the ∼1 and 50 mM NaCl conditions, FnCas12a ‘SKR’ and AsCas12a V1084K mutants had significantly better DNA detection than WT (**Figure 3D,E**). However, in more realistic sample matrix comprising 63% human saliva, these nucleases failed to generate a fluorescence signal different to the no-DNA control (**Figure 3F**). LbCas12a S929F/S1132R and WT both had very robust DNA detection across the ∼1 and 50 mM NaCl conditions, and could detect DNA spiked into the human saliva sample, with the LbS929F/S1132R mutant generating higher fluorescence (**Figure 3D, E, F**). This mutant had increased activity in higher NaCl concentrations (**Figure 3C**), which may account for its improved activity in electrolyte-rich human saliva (31).

We also assessed the gene editing activity of high-*trans* cleavage variants. Although AsV1084K showed only a modest increase in *trans* cleavage compared to WT, it had similar or higher gene editing activity across all target sites and cell lines tested (**Figure A,C,E**). However, it also had higher off-target activity when using the DNMT1-3 crRNA (**Figure S32,33**). This ∼1-3% off-target editing was low in comparison to the engineered AsCas12a ‘Ultra’ variant, which had up to 70% indels at this off-target site (**Figure S35**). Similar to a previous engineered Cas12a variant – enAsCas12a – AsCas12a ‘Ultra’ has increased activity at both on- and off-target sites (**Figure S34,35**) (27). It would appear mutations that allow wider PAM motif recognition also stabilises some degree of off-target activity. This increase is generally modest and can be mitigated by careful dosing of RNP in gene editing experiments (28). The increase in *k*_TS_, *trans* cleavage, and off-target editing by AsV1084K would suggest this mutation non-specifically stabilises the binding and cleavage of DNA strands.

When the *trans* cleavage of Cas12a was first characterised, there was some concern this indiscriminate DNAse activity may be deleterious in living cells (2). However, it has been demonstrated that the *trans* cleavage of AsCas12a or LbCas12a causes no detectable off-targets in mouse embryos (32). Similarly, *trans* cleavage had no discernible effect on plasmid or phage interference in *E. coli* (33, 34). This is likely due to the protection of ssDNA by binding proteins (35), higher ionic strength (36), and lower available amounts of Mg^2+^ ions in living cells compared to *in vitro* (37). Thus, the AsCas12a V1084K variant is unlikely to cause unwanted off-target genome editing through enhanced *trans* cleavage, instead it is likely through stabilising off-target DNA binding and *cis* cleavage.

For example, the high-*trans* LbCas12a S929F/S1132R mutant displayed similar gene-editing activity to WT, and significantly higher in one instance (**Figure 4B,D,F**), with consistently low off-target activity (**Figure S31-33**). This modest effect would suggest these mutations make a minor difference in living cells.

### Implications for high- and low-*trans cleaving Cas12a variants*

With an explosion of discovery and characterisation of a variety of Cas12a orthologues, and their use in molecular detection and genome editing (5, 38, 39), the ability to modulate *cis* and *trans* cleavage may be widely useful. For example, recent engineering of the NTS-binding groove of a thermostable Cas12a orthologue was able to increase its *trans* cleavage, for more sensitive RT-LAMP enabled detection of RNA (16). Further engineering of the Nuc and RuvC-lid of this orthologue has the potential to further enhance its activity.

Although *trans* cleavage is central to generating a signal in Cas12a molecular detection, recent work has emphasised the importance of enzyme activation kinetics (40). Before *cis* or *trans* cleavage, a Cas12a-crRNA binary complex must locate the target sequence by non-specific DNA diffusion and specific PAM motif binding (41). This step is rate-limiting in detecting low-abundance DNA targets, and the more rapid activation kinetics of the TsCas12a orthologue generated a signal faster than LbCas12a, despite higher steady-state *trans* cleavage for the latter (40). Orthologue-informed mutation of the Nuc domain of TsCas12a may enhance its steady-state *trans* cleavage, for more sensitive room-temperature diagnostics (40).

Mutations that decreased *trans* cleavage could also decrease *cis* cleavage. This is likely by decreasing the strength of nonspecific protein-nucleic acid interactions near the RuvC. This could be leveraged to improve the gene editing specificity of Cas12a orthologues. The specificity of Cas12a has been previously improved by weakening interactions between the REC lobe and the crRNA-TS heteroduplex (27). The effect of alanine substitution mutants in the Nuc domain could be explored in a similar fashion. Although our triple and quadruple alanine substitution mutants of AsCas12a and LbCas12a had overall lower on-target gene editing activity (**Figure 4**), it is possible that single or double mutants could retain more on-target activity.

## Conclusions

In this work, we studied the role of the residues near the RuvC on the *cis* and *trans* DNA cleavage activities of three Cas12a orthologues (**Figure 5**). We found that alanine substitution of arginine or lysine residues in the Nuc domain can abolish the *trans* cleavage of Cas12a orthologues, while modestly reducing *cis* cleavage rates. We replicated findings in the literature that increasing NaCl concentration in reaction buffers decreases *trans* cleavage rates, and suggest these ions neutralise electrostatic interactions between positively charged Nuc residues and the negatively charged phosphate backbone of *trans* substrates. As these electrostatic interactions are critical, substituting additional arginine or lysine residues in the Nuc was able to increase *trans* cleavage rates, especially at higher ionic strengths. Substitutions of the RuvC-lid could also increase *trans* cleavage, but not predictably. Testing combinations of RuvC-lid and Nuc mutations yielded high- and low-*trans* cleaving variants of three Cas12 orthologues. This study provides a blueprint for future rational engineering of Cas12a nucleases for their *trans* cleveage activities.

## Acknowledgements

The authors would like to thank the Biomolecular Resource Facility at the ANU for undertaking the DNA sequencing.

## Author contributions

A.N. performed structural analyses, plasmid cloning and protein purification, and assays for protein thermostability, *cis* and *trans* cleavage, and DNA detection. L.S. performed human cell culturing, electroporations, and genomic DNA extractions. A.N. and G.B. analysed *cis* and *trans* cleavage rates, and high-throughput sequencing data. A.N. and G.B conceived the study and wrote the manuscript. All authors commented and approved the manuscript.

## Supplementary Data

Available in attached file.

## Data availability

Data from high-throughput sequencing have been deposited with the National Center for Biotechnology Information Sequence Read Archive under BioProject ID PRJNA1281281. All other datasets are available on request.

