## Supplementary data for "Nuc domain electrostatics drive the *trans* cleavage activity of CRISPR-Cas12a"

**for**

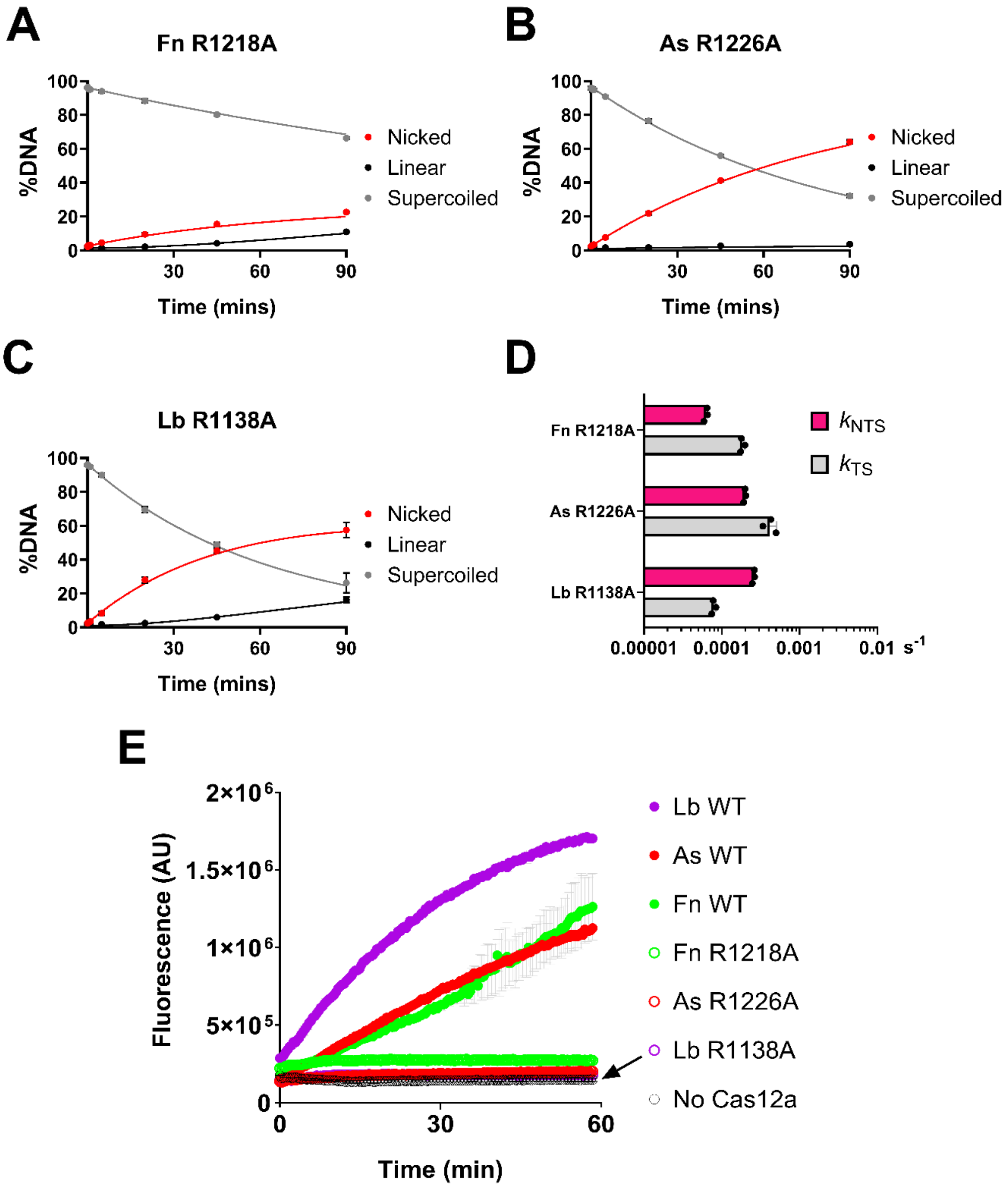


**Figure S1:** Time-course plasmid *cis* cleavage by (**A**) FnR1218A, (**B**), AsR1226A, and (**C**) LbR1138A. Dots show mean, error bars show s.d. Lines show fit of sequential NTS and TS cleavage model. (**D**) Kinetic values ($s^{-1}$) of NTS ($k_{NTS}$, pink) and TS cleavage ($k_{TS}$, grey), for the Cas12a indicated. Dots show individual replicates, bars show mean, error bars show s.d. (**E**) *Trans* cleavage of WT and mutants of FnCas12a, AsCas12a, and LbCas12a, when activated by 1 nM DNMT1 target dsDNA. Dots show mean, error bars show s.d.

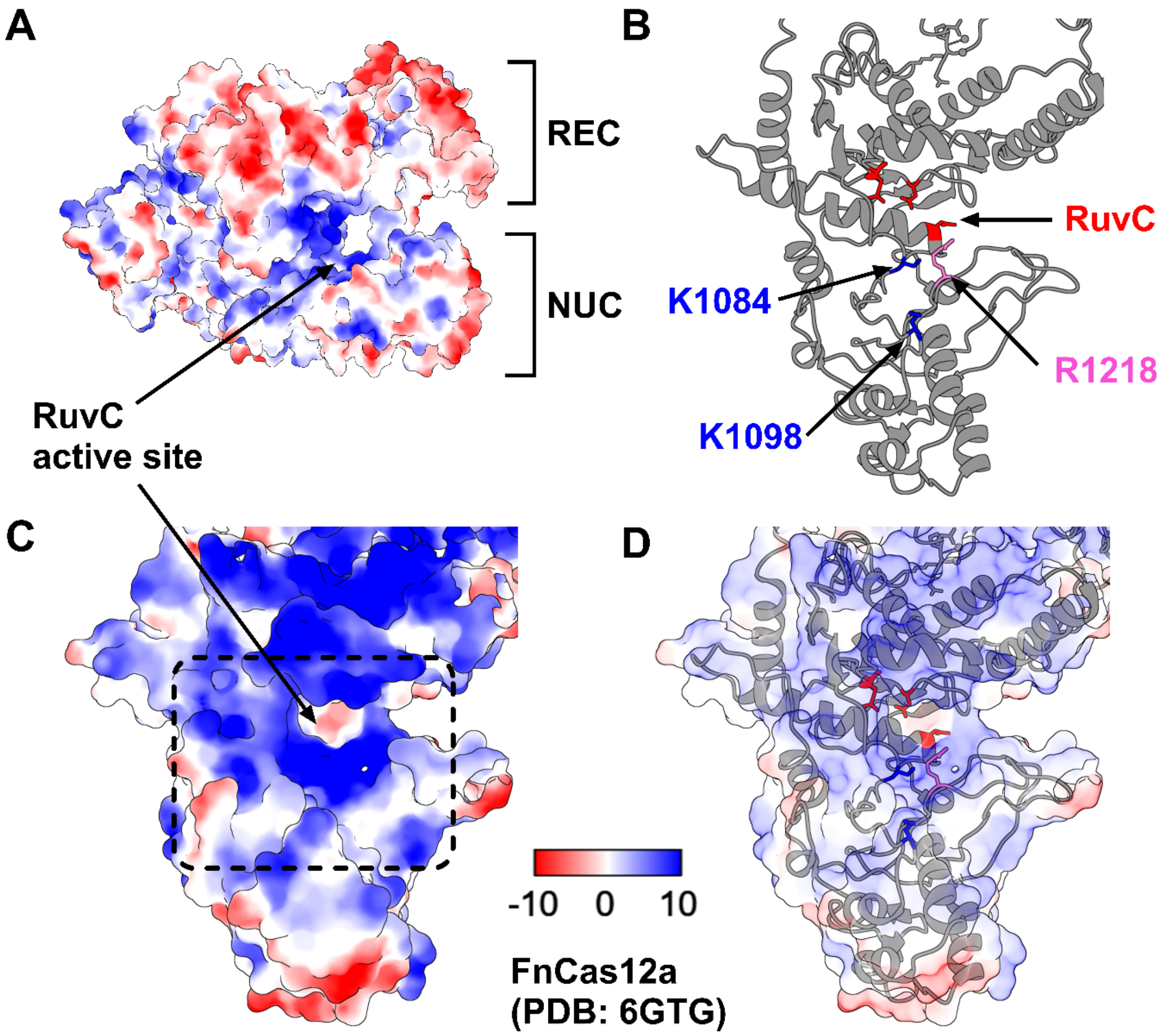


**FigureS2:** (**A**) Coulombic electrostatic potential of FnCas12a (PDB: 6GTG), as calculated by ChimeraX. Electrostatic potential is displayed on a colour gradient showing negative (red), neutral (white), and positive (blue). REC and NUC lobes are annotated, as is location of RuvC active site. (**B**) Ribbon representation of the NUC lobe, with key amino acids highlighted; RuvC active site residues (red), conserved adjacent arginine (pink), and positively charged residues subject to alanine substitution (blue). (**C**) Electrostatic potential of the NUC lobe, with arrow showing RuvC site. Dashed box shows cropped region shown in **Figure 1A** of main text. (**D**) Merge of ribbon representation with electrostatic potential (at 80% transparency).

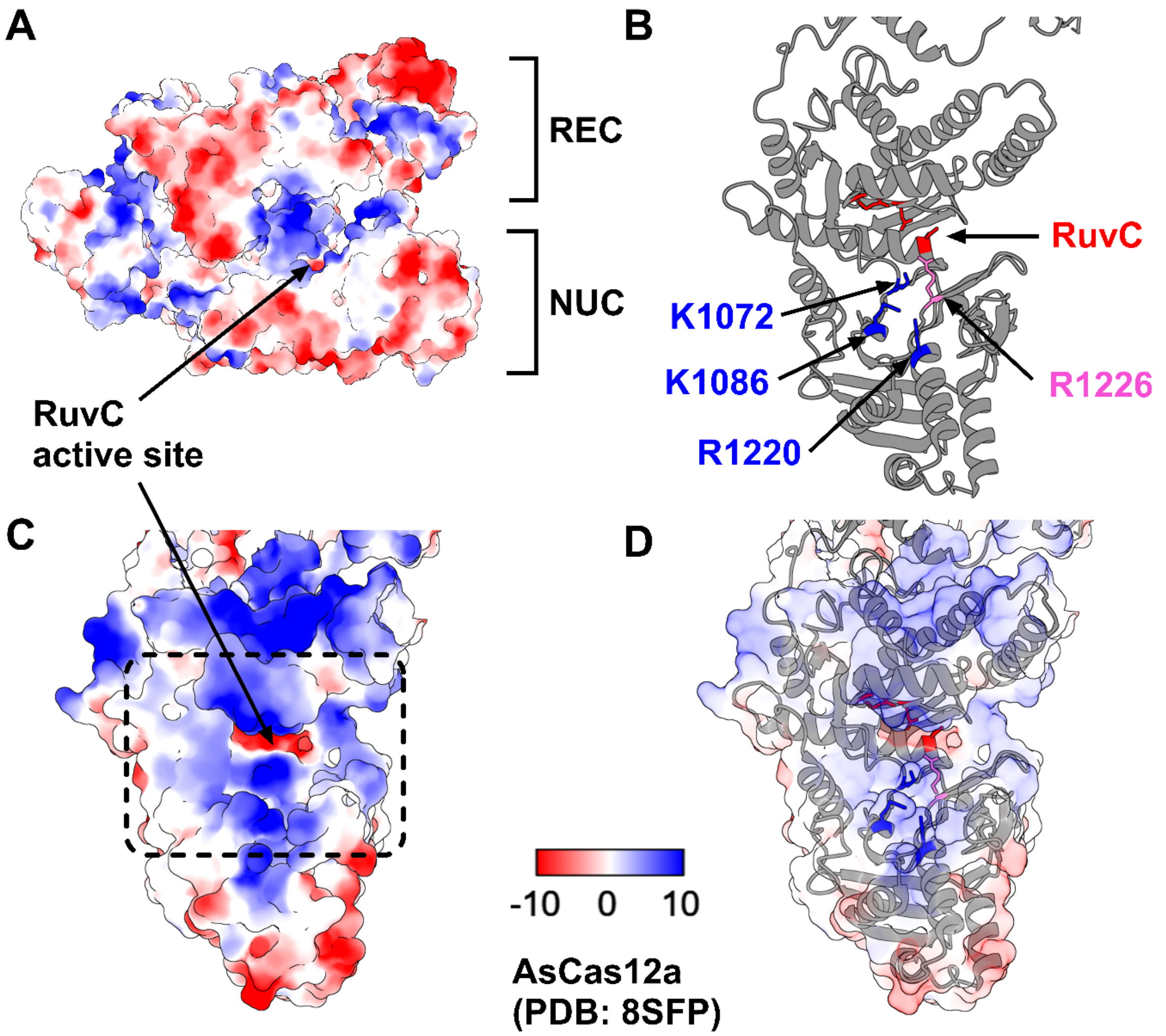


**FigureS3:** (**A**) Coulombic electrostatic potential of AsCas12a (PDB: 8SFP), as calculated by ChimeraX. Electrostatic potential is displayed on a colour gradient showing negative (red), neutral (white), and positive (blue). REC and NUC lobes are annotated, as is location of RuvC active site. (**B**) Ribbon representation of the NUC lobe, with key amino acids highlighted; RuvC active site residues (red), conserved adjacent arginine (pink), and positively charged residues subject to alanine substitution (blue). (**C**) Electrostatic potential of the NUC lobe, with arrow showing RuvC site. Dashed box shows cropped region shown in **Figure 1B** of main text. (**D**) Merge of ribbon representation with electrostatic potential (at 80% transparency).

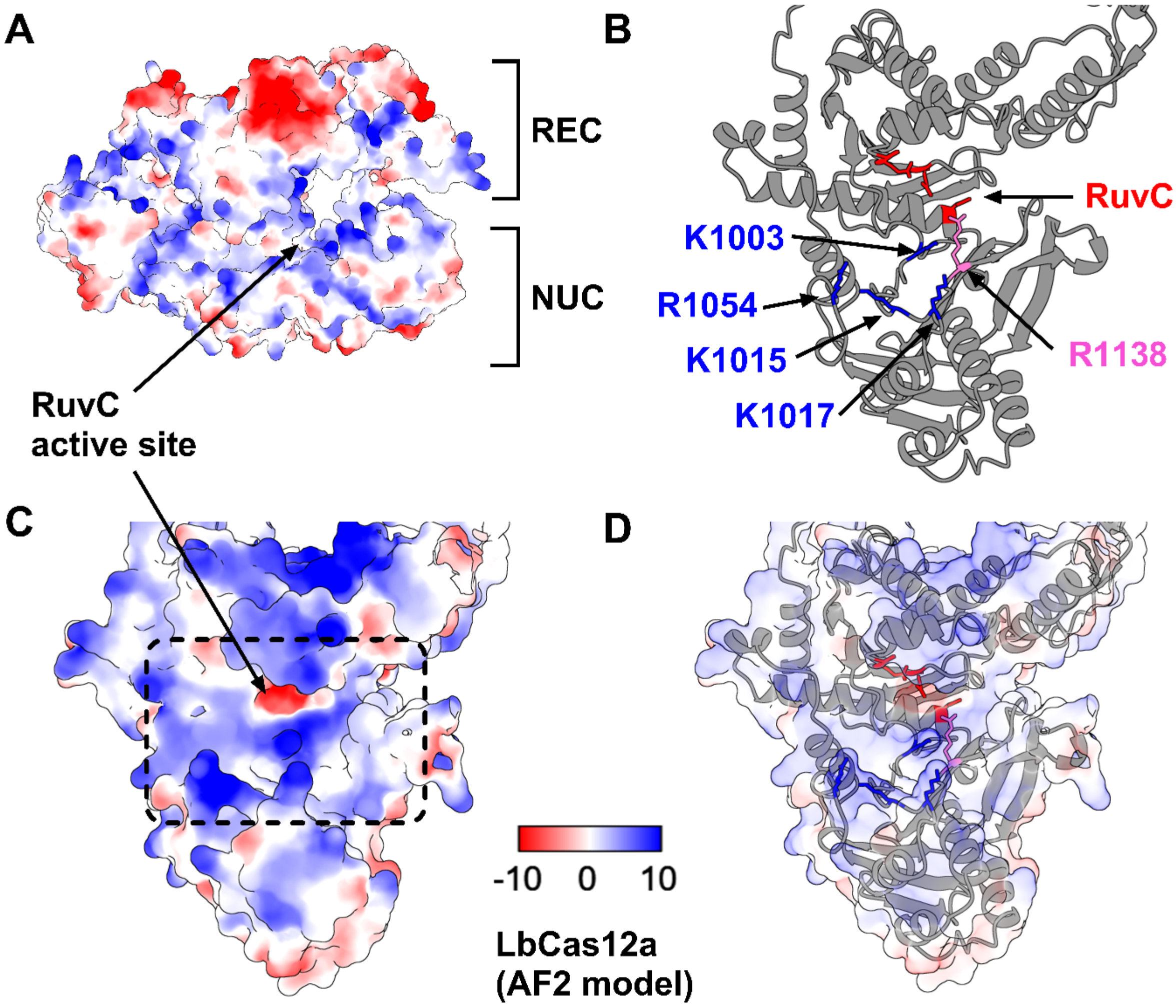


**FigureS4:** (**A**) Coulombic electrostatic potential of LbCas12a (AlphaFold monomer V2.0 prediction of A0A182DWE3), as calculated by ChimeraX. Electrostatic potential is displayed on a colour gradient showing negative (red), neutral (white), and positive (blue). REC and NUC lobes are annotated, as is location of RuvC active site. (**B**) Ribbon representation of the NUC lobe, with key amino acids highlighted; RuvC active site residues (red), conserved adjacent arginine (pink), and positively charged residues subject to alanine substitution (blue). (**C**) Electrostatic potential of the NUC lobe, with arrow showing RuvC site. Dashed box shows cropped region shown in **Figure 1C** of main text. (**D**) Merge of ribbon representation with electrostatic potential (at 80% transparency).

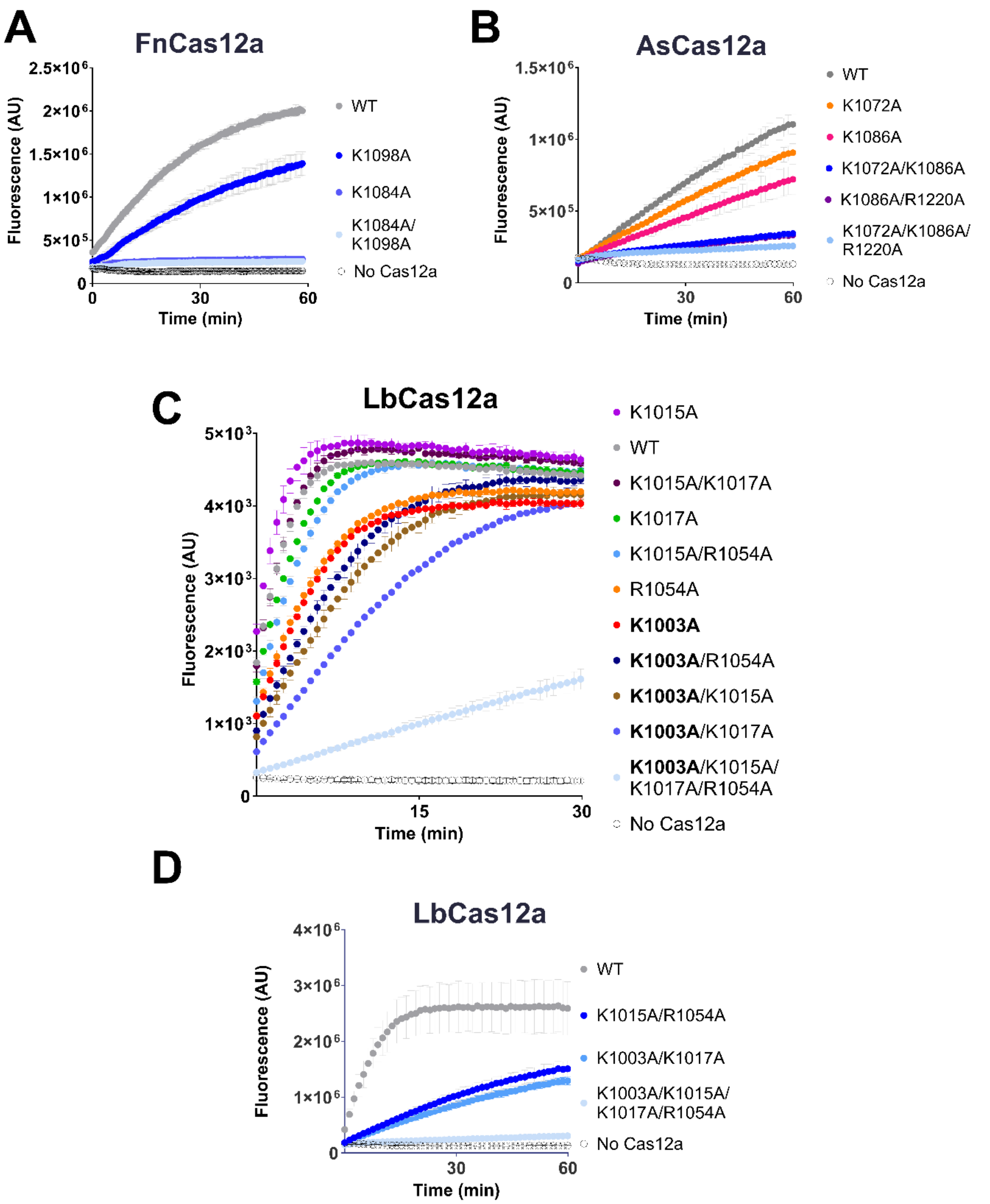


**Figure S5:** (**A**) *Trans* cleavage curve of WT FnCas12a and mutants, when activated by 1 nM DNMT1 dsDNA. (**B**) *Trans* cleavage curve of WT AsCas12a and mutants when activated by 1 nM DNMT1 dsDNA. (**C**) *Trans* cleavage curve of WT LbCas12a and mutants when activated by 5 nM DNMT1 target-strand ssDNA. Key mutation K1003A in bold. (**D**) *Trans* cleavage curve of WT LbCas12a and mutants, when activated by 1 nM DNMT1 dsDNA. Reaction buffers contained 50 mM NaCl (final concentration).

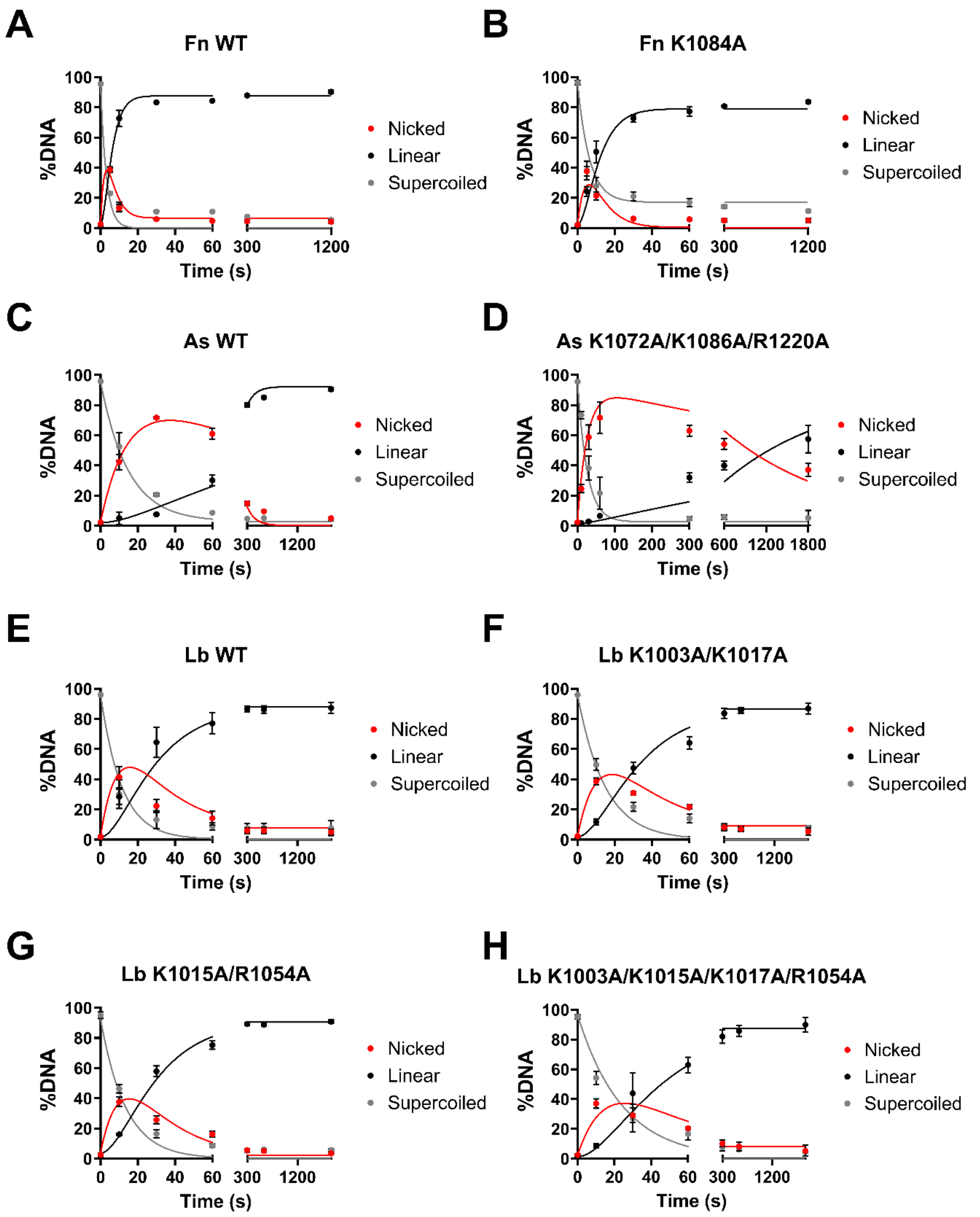


**Figure S6:** Time-course plasmid *cis* cleavage by (**A**) FnCas12a WT, (**B**) FnK1084A, (**C**) AsCas12a WT, (**D**) AsK1072A/K1086A/R1220A, (**E**) LbCas12a WT, (**F**) LbK1003A/K1017A, (**G**) LbK1015A/R1054A, (**H**) LbK1003A/K1015A/K1017A/R1054A. Dots show mean, error bars show s.d. Lines show fit of sequential NTS and TS cleavage model. Conducted in reaction buffers with a final concentration of 50 mM NaCl.

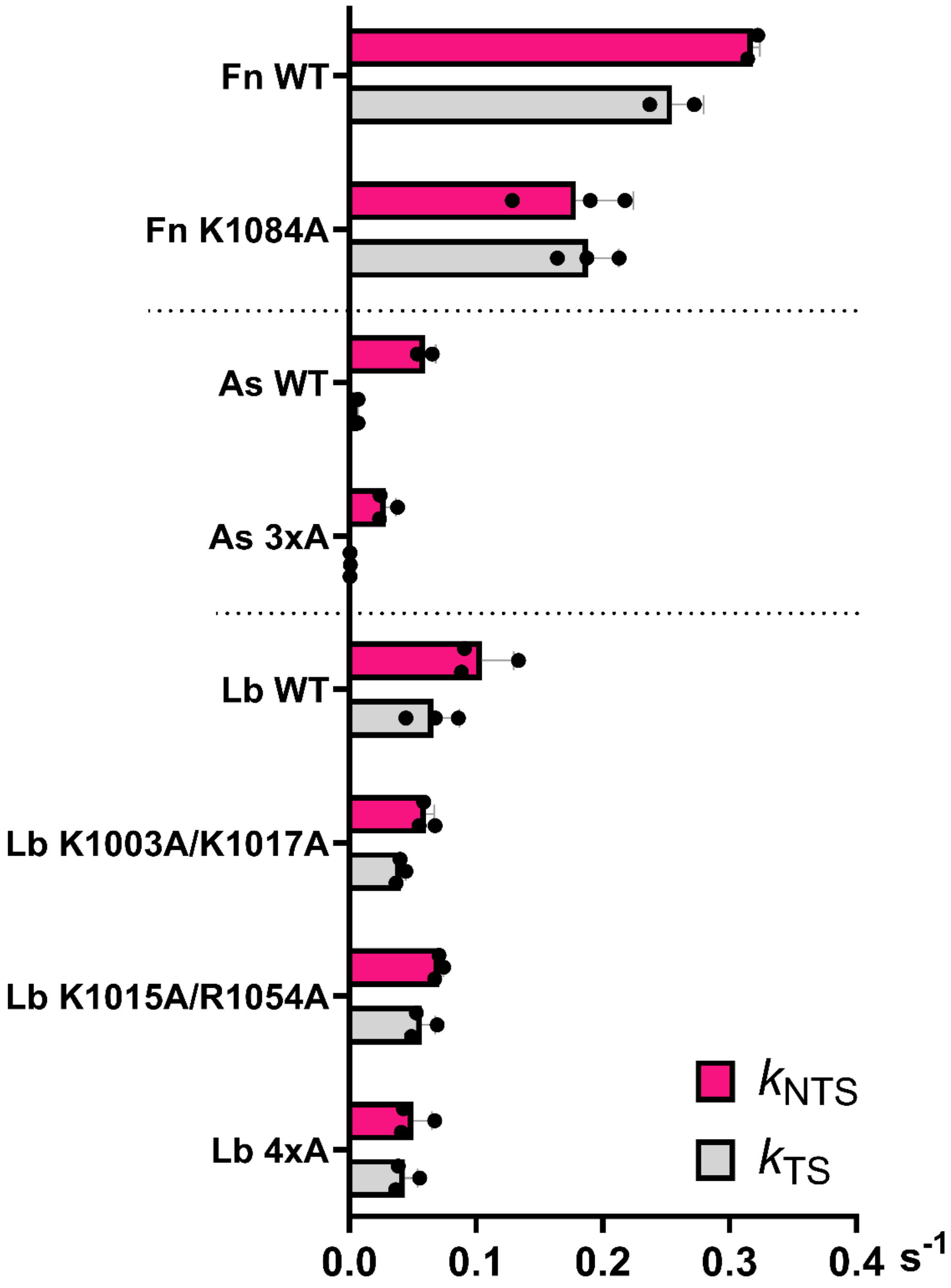


**Figure S7:** Kinetic values ($s^{-1}$) of NTS ($k_{NTS}$, pink) and TS cleavage ($k_{TS}$, grey), for the Cas12a indicated. Dots show individual replicates, bars show mean, error bars show s.d. Derived from **Figure S6** for the Cas12a indicated.

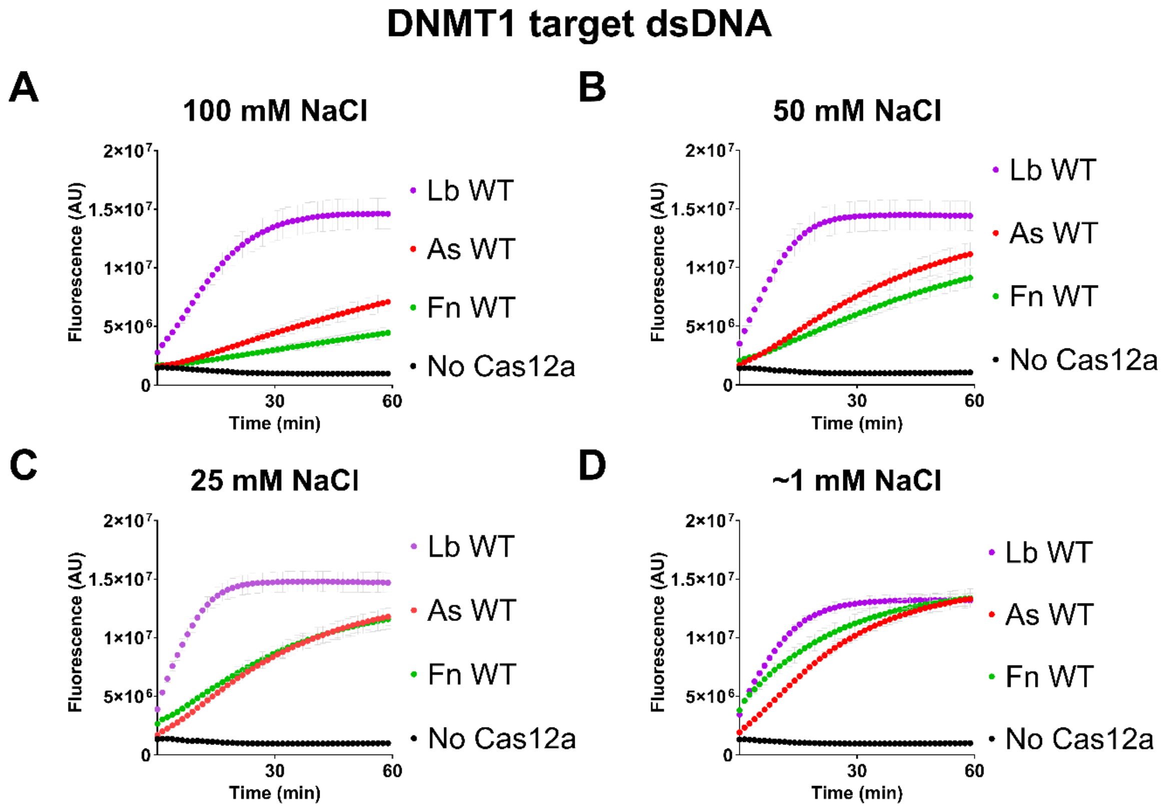


**Figure S8**: *Trans* cleavage curves of WT Cas12a orthologues, when activated by 1 nM DNMT1 dsDNA, in reaction buffers containing a final NaCl concentration of (**A**) 100 mM, (**B**) 50 mM, (**C**) 25 mM, (**D**) ~1 mM.

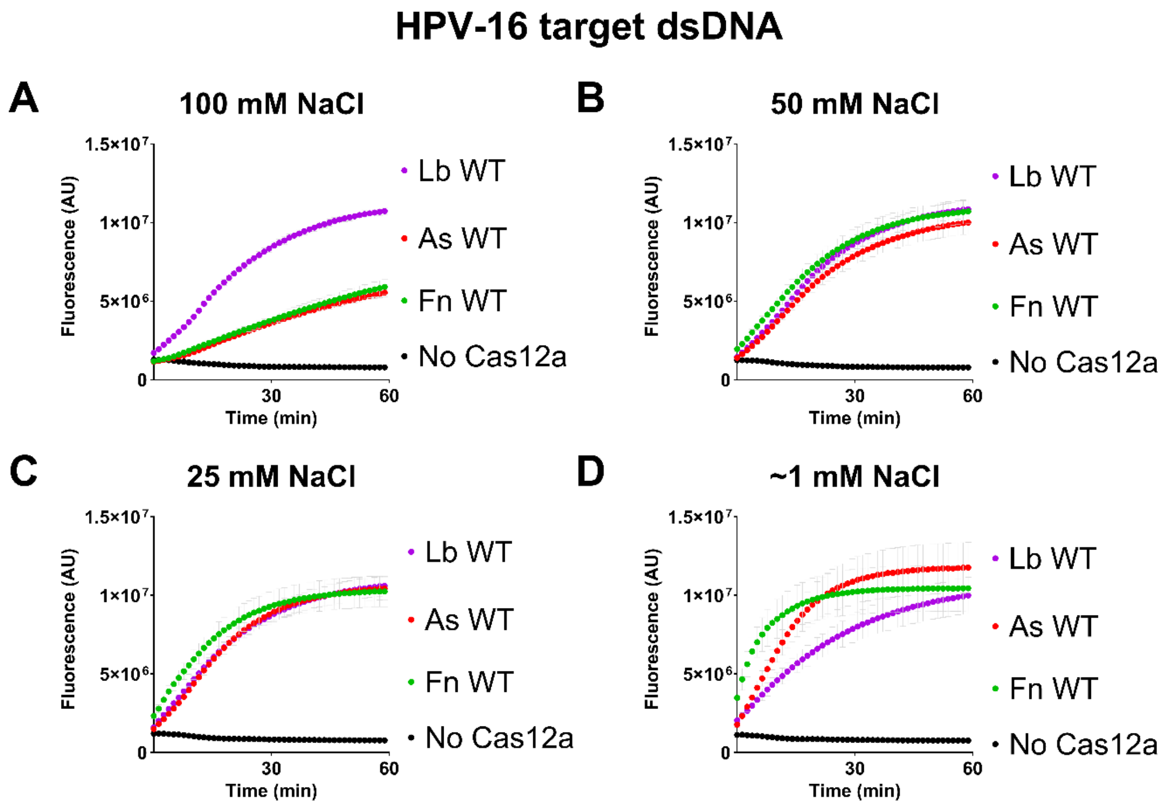


**Figure S9:** *Trans* cleavage curves of WT Cas12a orthologues, when activated by 1 nM HPV-16 dsDNA, in reaction buffers containing a final NaCl concentration of (**A**) 100 mM, (**B**) 50 mM, (**C**) 25 mM, (**D**) ~1 mM.

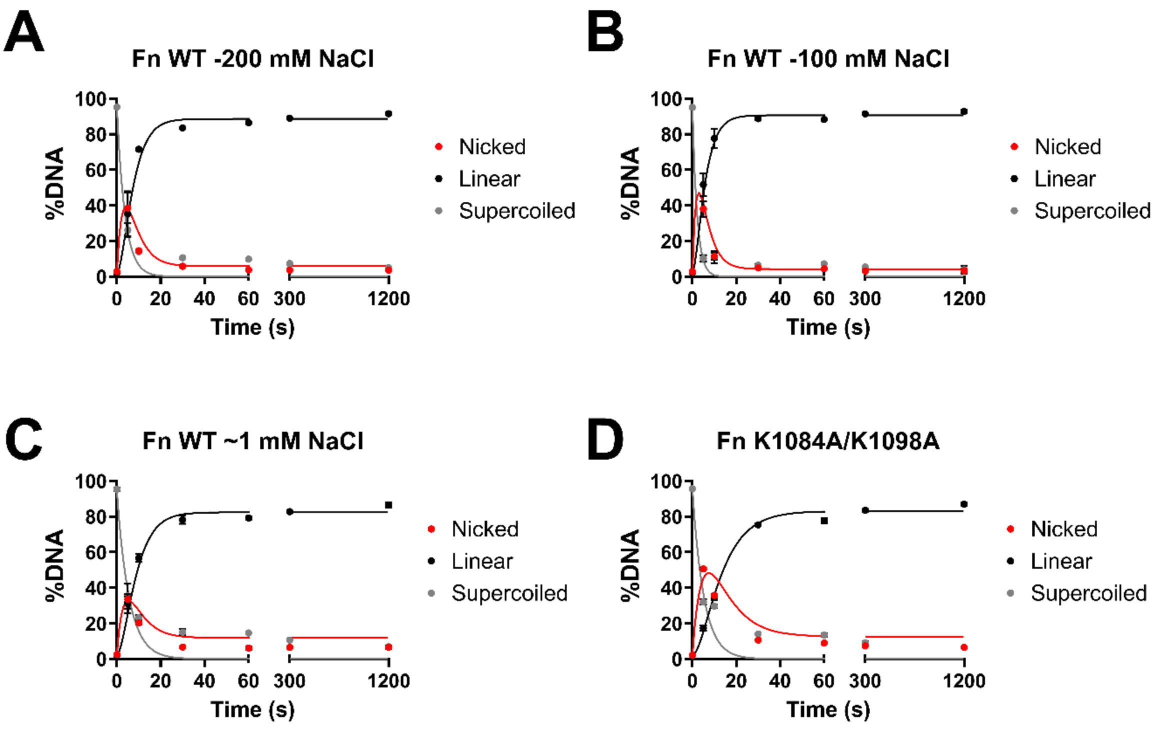


**Figure S10:** Time-course plasmid *cis* cleavage of WT FnCas12a, in reaction buffers containing a final NaCl concentration of (**A**) 200 mM, (**B**) 100 mM, (**C**) ~1 mM. (**D**) *Cis* cleavage kinetics of FnCas12a K1084A/K1098A mutant, in a final concentration of 50 mM NaCl. Dots show mean, error bars show s.d. Lines show fit of sequential NTS and TS cleavage model.

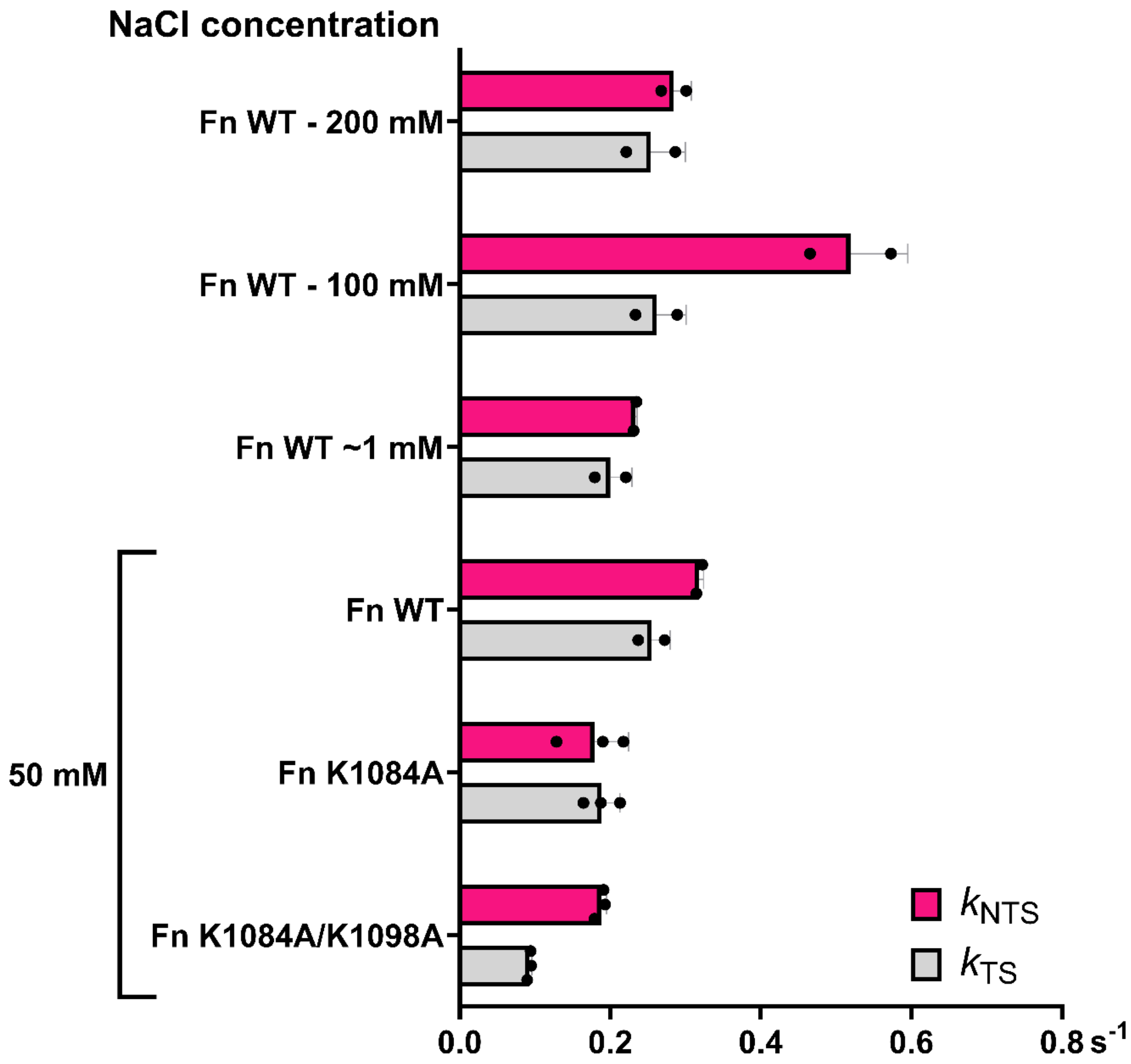


**Figure S11:** Kinetic values ($s^{-1}$) of NTS ($k_{NTS}$, pink) and TS cleavage ($k_{TS}$, grey), for the Cas12a indicated. Dots show individual replicates, bars show mean, error bars show s.d. Derived from **Figure S6A** and **Figure S10** for the sample indicated.

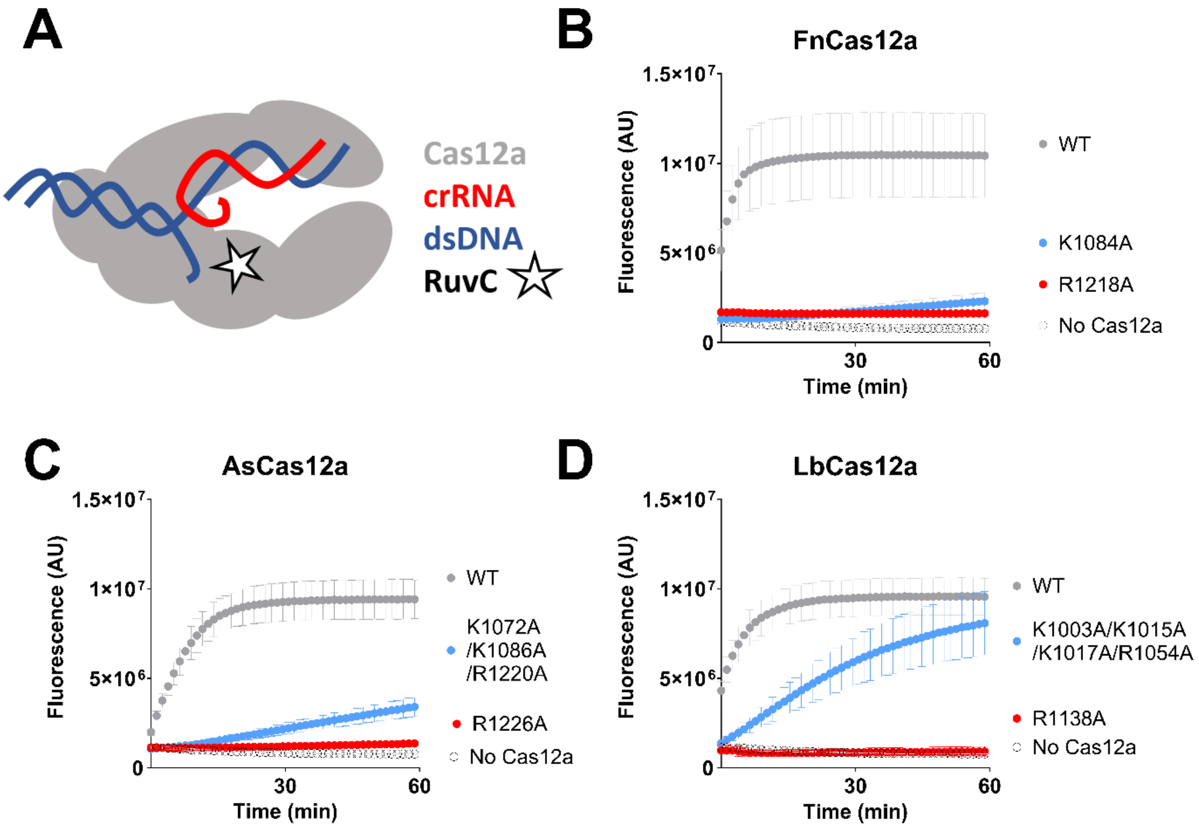


**Figure S12:** (**A**) Diagram of Cas12a bound to a 'pre-cleaved' dsDNA substrate. *Trans* cleavage of (**B**) FnCas12a WT and mutants, (**C**) AsCas12a WT and mutants, (**D**) LbCas12a WT and mutants, when activated by 1 nM 'pre-cleaved' dsDNA substrate.

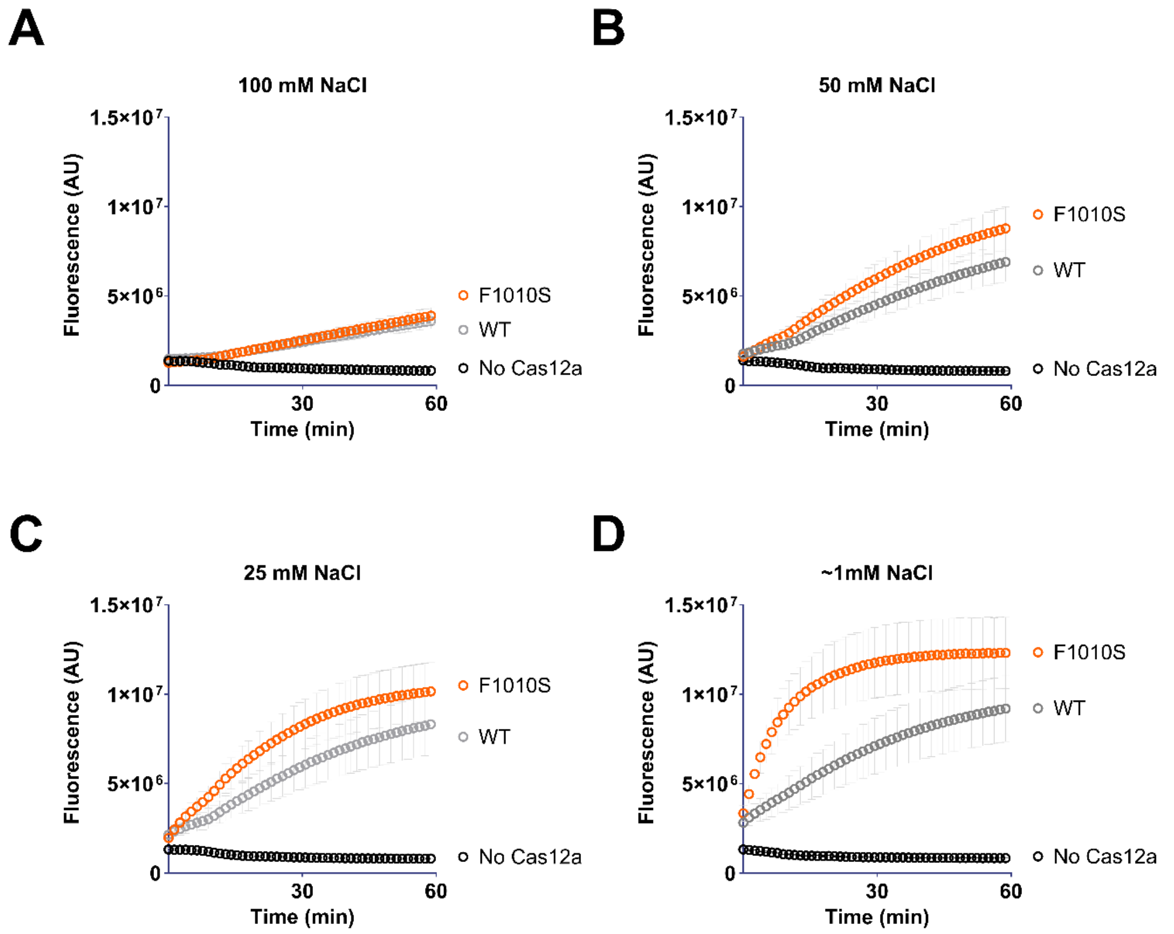


**FigureS13:** *Trans* cleavage curves of WT FnCas12a and F1010S mutant, when activated by 1 nM DNMT1 dsDNA, in reaction buffers containing a final NaCl concentration of (**A**) 100 mM, (**B**) 50 mM, (**C**) 25 mM, (**D**) ~1 mM.

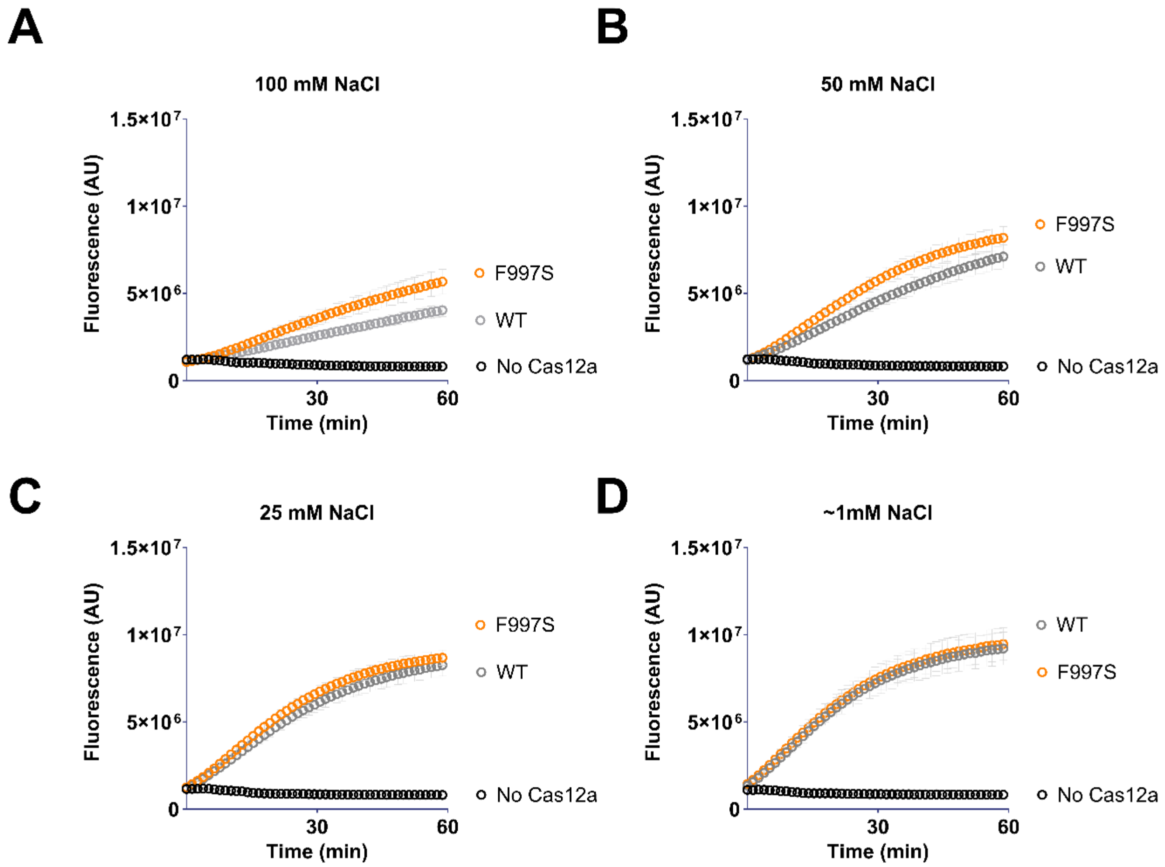


**FigureS14:** *Trans* cleavage curves of WT AsCas12a and F997S mutant, when activated by 1 nM DNMT1 dsDNA, in reaction buffers containing a final NaCl concentration of (**A**) 100 mM, (**B**) 50 mM, (**C**) 25 mM, (**D**) ~1 mM.

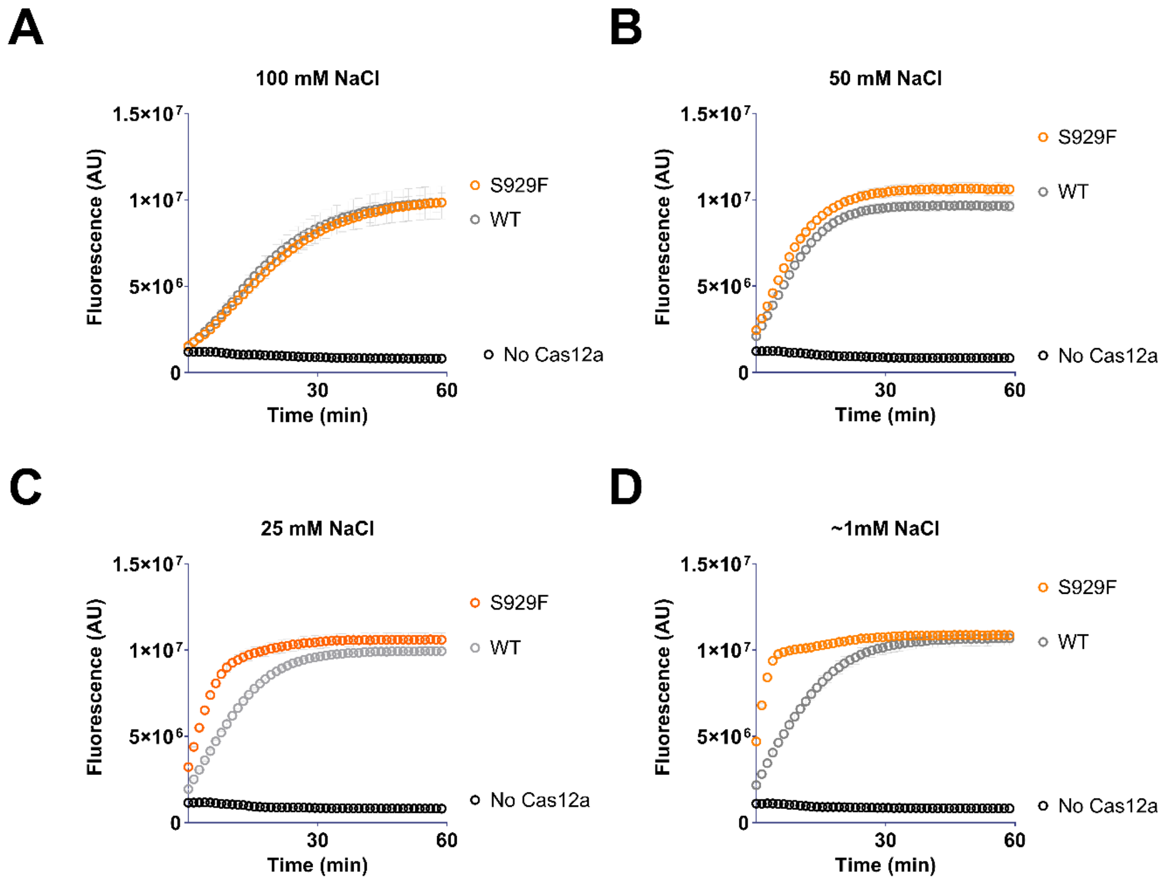


**FigureS15:** *Trans* cleavage curves of WT LbCas12a and S929F mutant, when activated by 1 nM DNMT1 dsDNA, in reaction buffers containing a final NaCl concentration of (**A**) 100 mM, (**B**) 50 mM, (**C**) 25 mM, (**D**) ~1 mM.

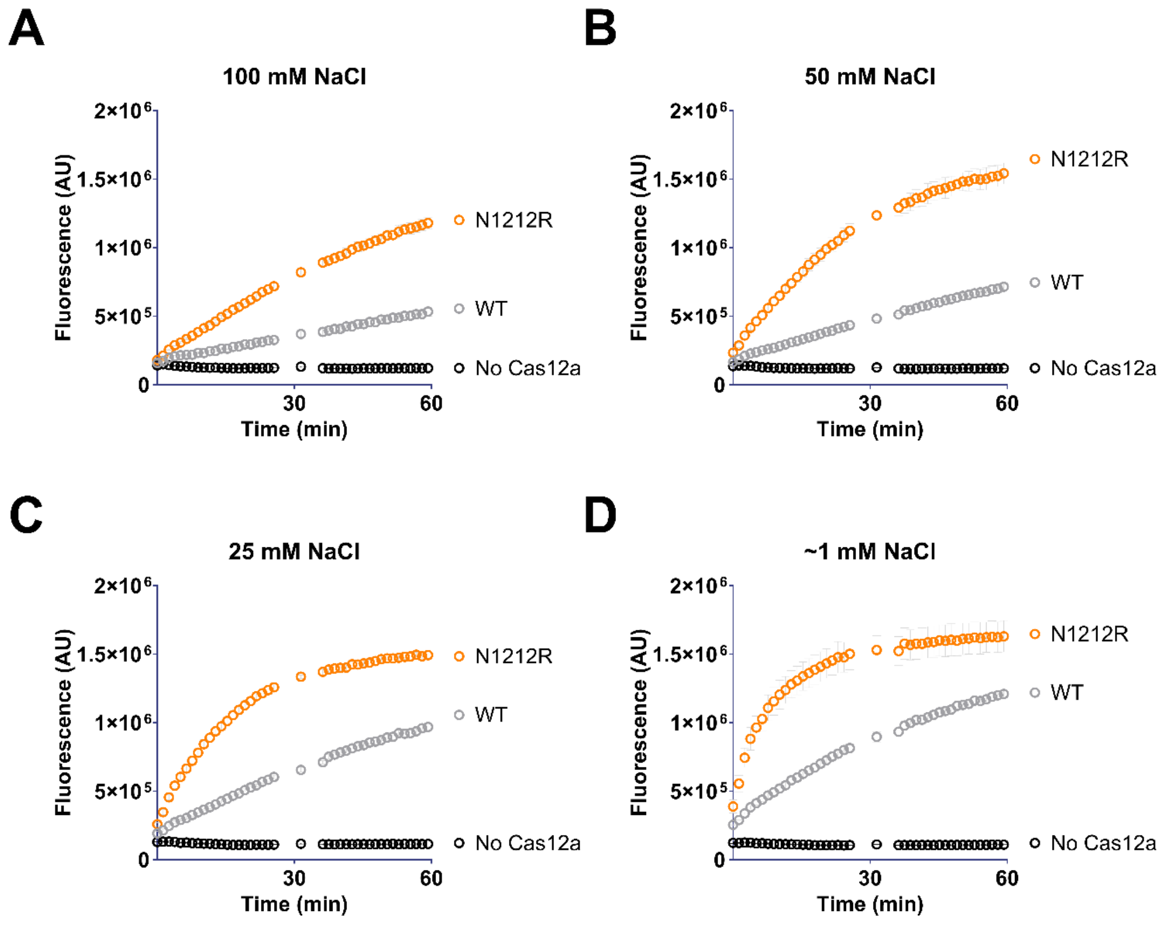


**Figure S16:** *Trans* cleavage curves of WT FnCas12a and N1212R mutant, when activated by 1 nM DNMT1 dsDNA, in reaction buffers containing a final NaCl concentration of (**A**) 100 mM, (**B**) 50 mM, (**C**) 25 mM, (**D**) ~1 mM. Missing data points due to machine error.

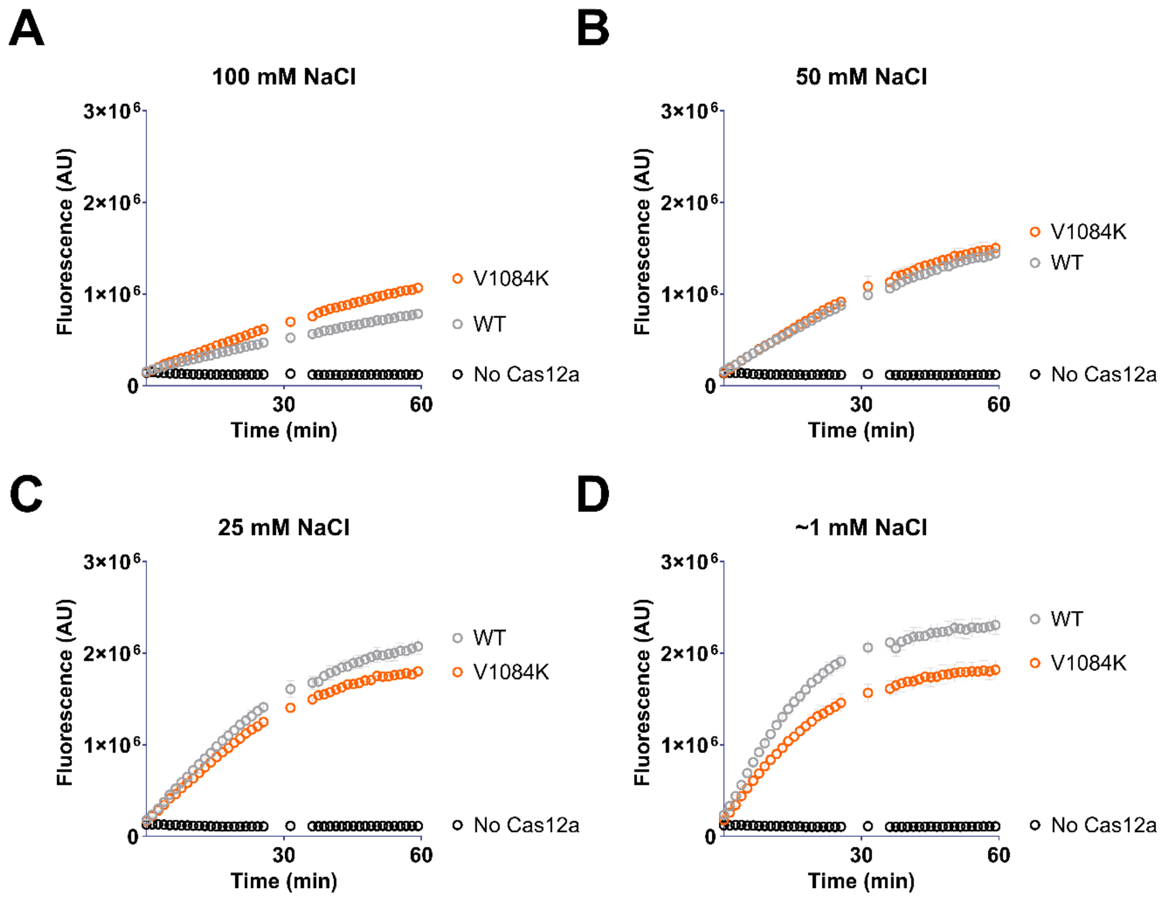


**Figure S17:** *Trans* cleavage curves of WT AsCas12a and V1084K mutant, when activated by 1 nM DNMT1 dsDNA, in reaction buffers containing a final NaCl concentration of (**A**) 100 mM, (**B**) 50 mM, (**C**) 25 mM, (**D**) ~1 mM. Missing data points due to machine error.

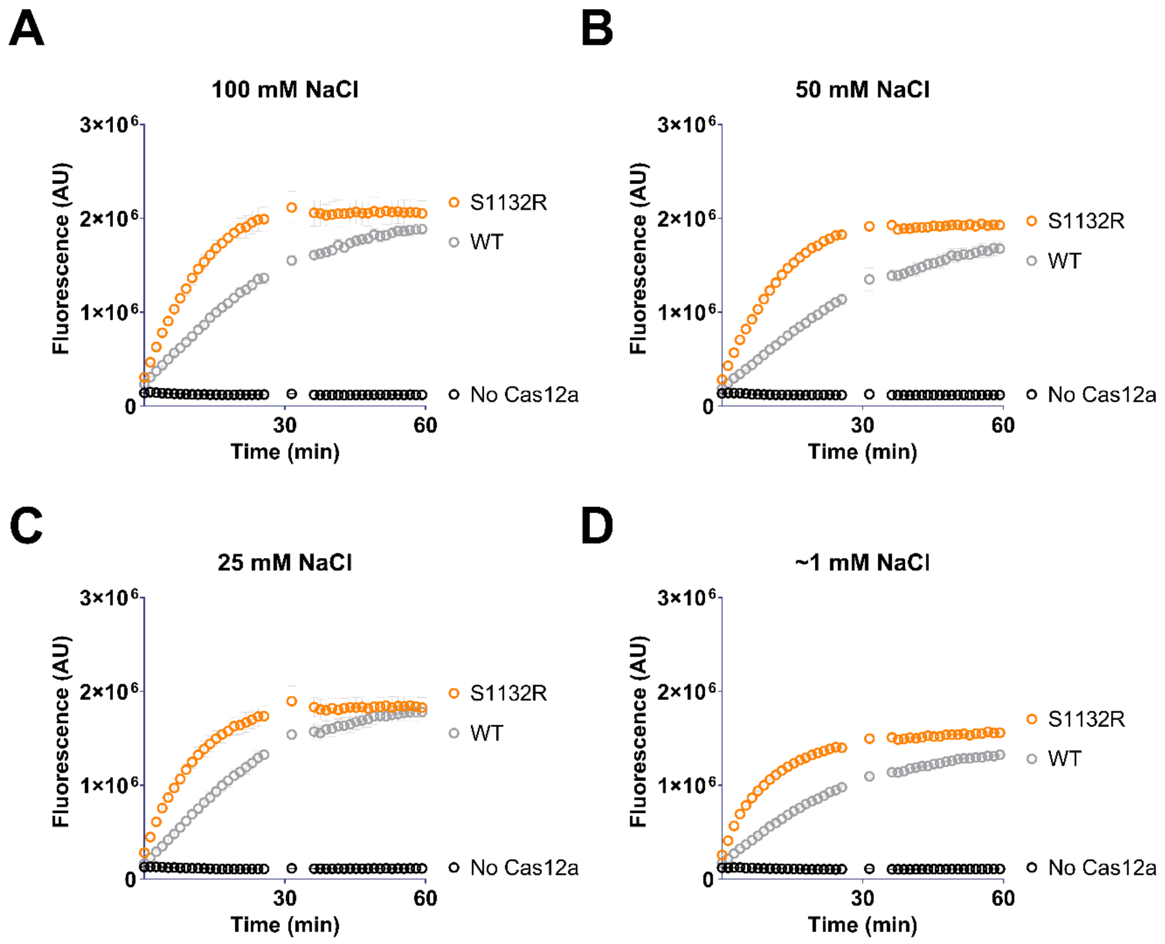


**Figure S18:** *Trans* cleavage curves of WT LbCas12a and S1132R mutant, when activated by 1 nM DNMT1 dsDNA, in reaction buffers containing a final NaCl concentration of (**A**) 100 mM, (**B**) 50 mM, (**C**) 25 mM, (**D**) ~1 mM. Missing data points due to machine error.

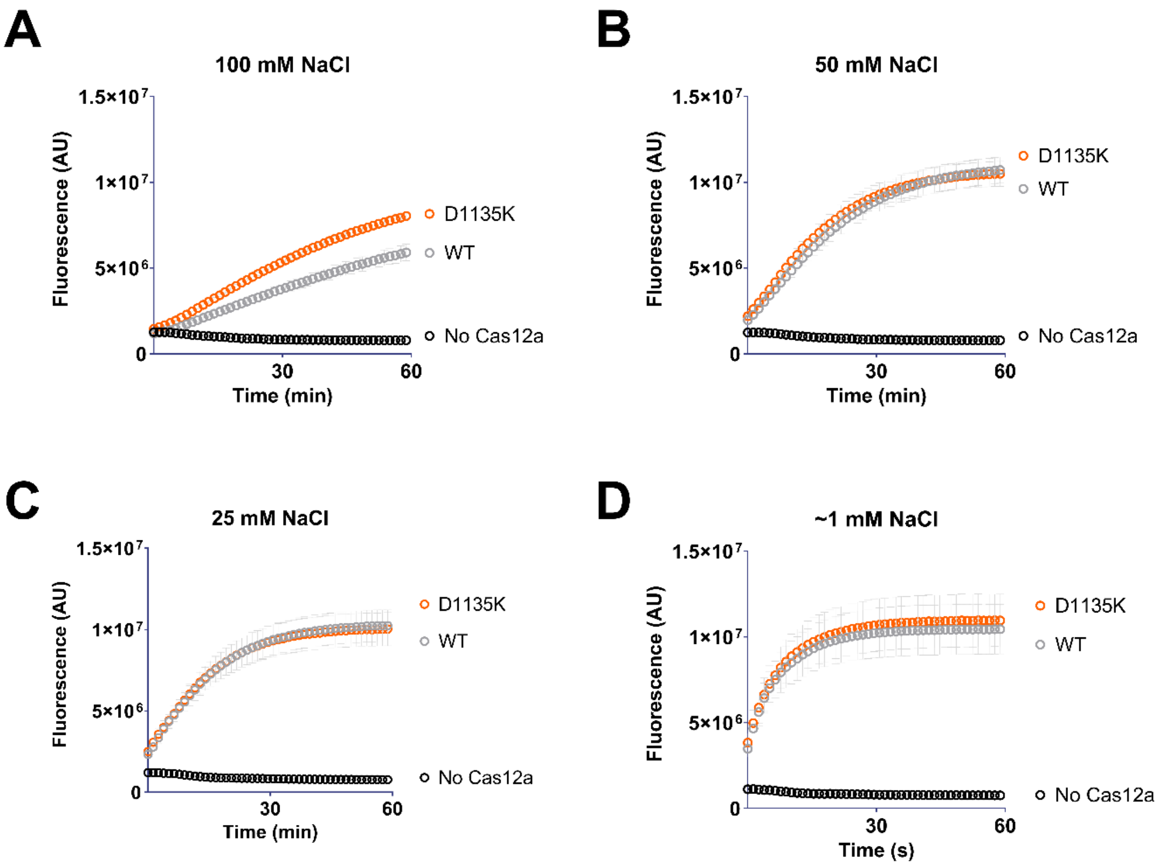


**Figure S19:** *Trans* cleavage curves of WT FnCas12a, and D1135K mutant, when activated by 1 nM DNMT1 dsDNA, in reaction buffers containing a final NaCl concentration of (**A**) 100 mM, (**B**) 50 mM, (**C**) 25 mM, (**D**) ~1 mM.

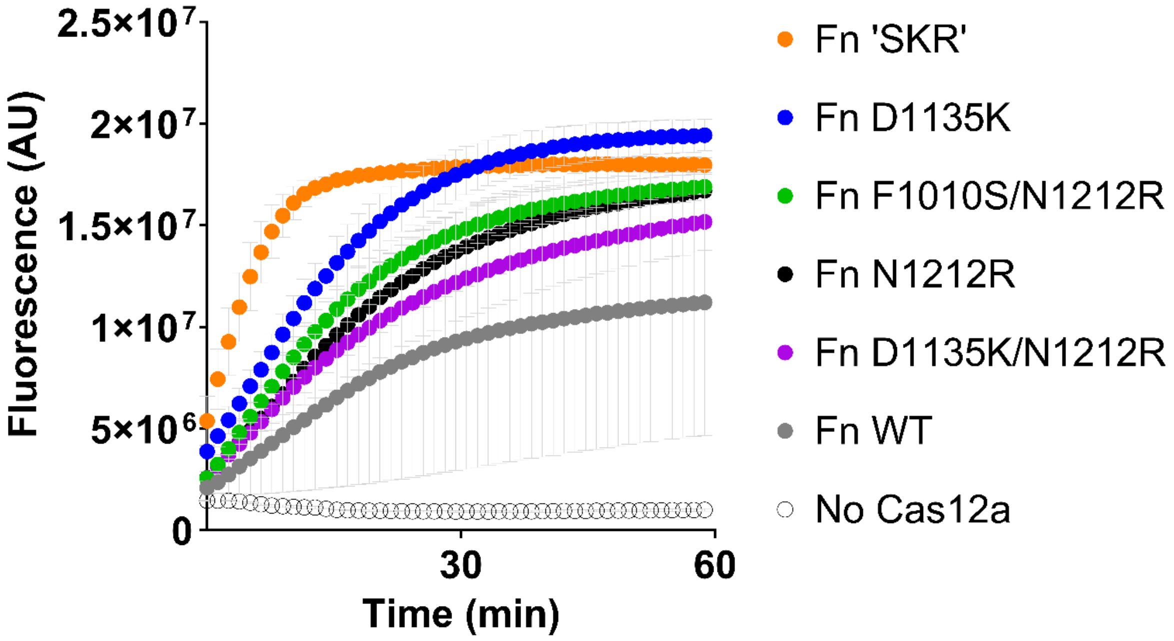


**Figure S20:** *Trans* cleavage curves of FnCas12a WT and mutants (as indicated), when activated by 1 nM DNMT1 dsDNA, in reaction buffer containing 50 mM NaCl.

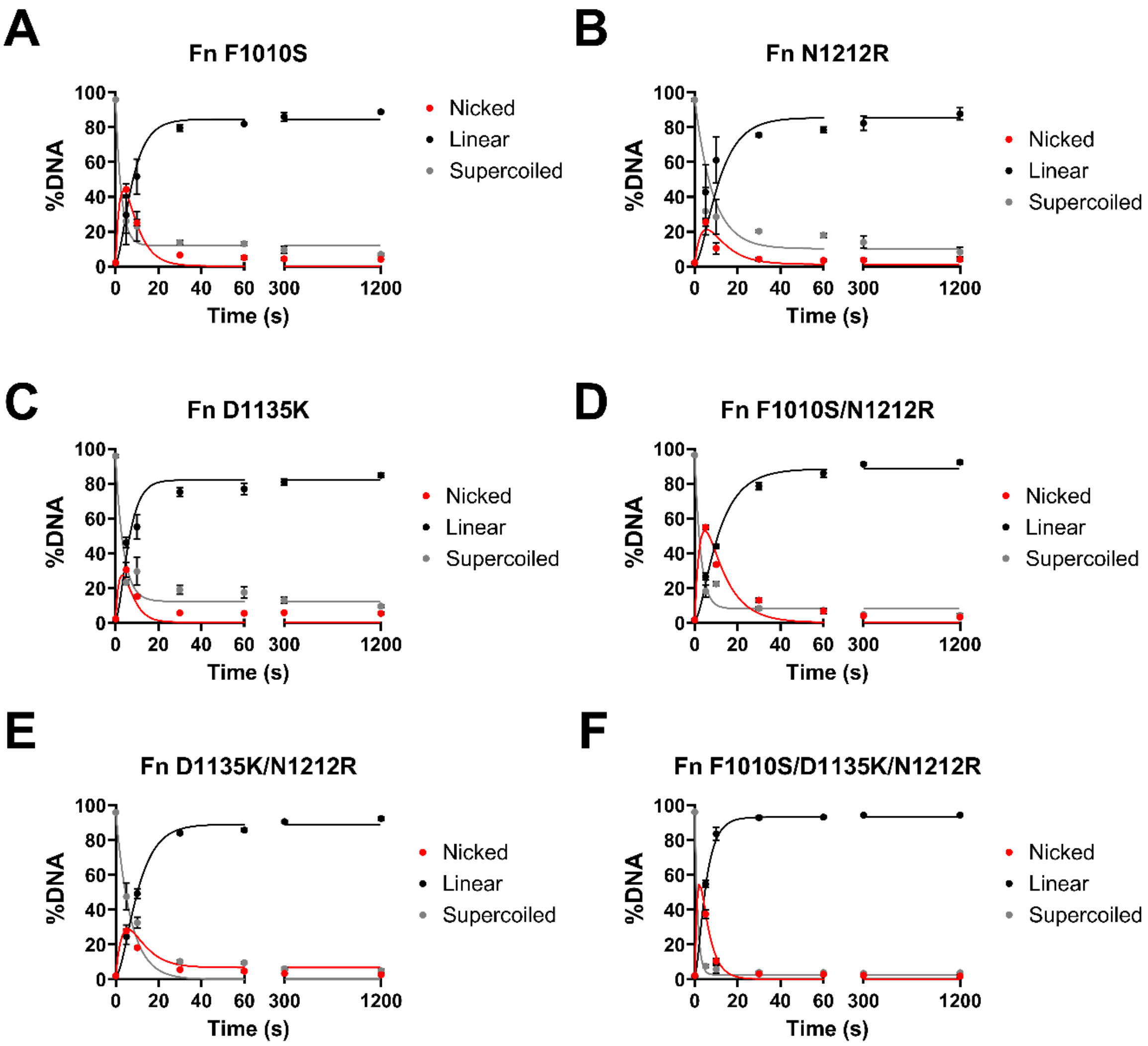


**Figure S21:** Time-course plasmid *cis* cleavage by (**A**) FnF1010S, (**B**) FnN1212R, (**C**) FnD1135K, (**D**) FnF1010S/N1212R, (**E**) FnD1135K/N1212R, and (**F**) FnF1010S/D1135K/N1212R. Dots show mean, error bars show s.d. Lines show fit of sequential NTS and TS cleavage model. Conducted in reaction buffers with a final concentration of 50 mM NaCl.

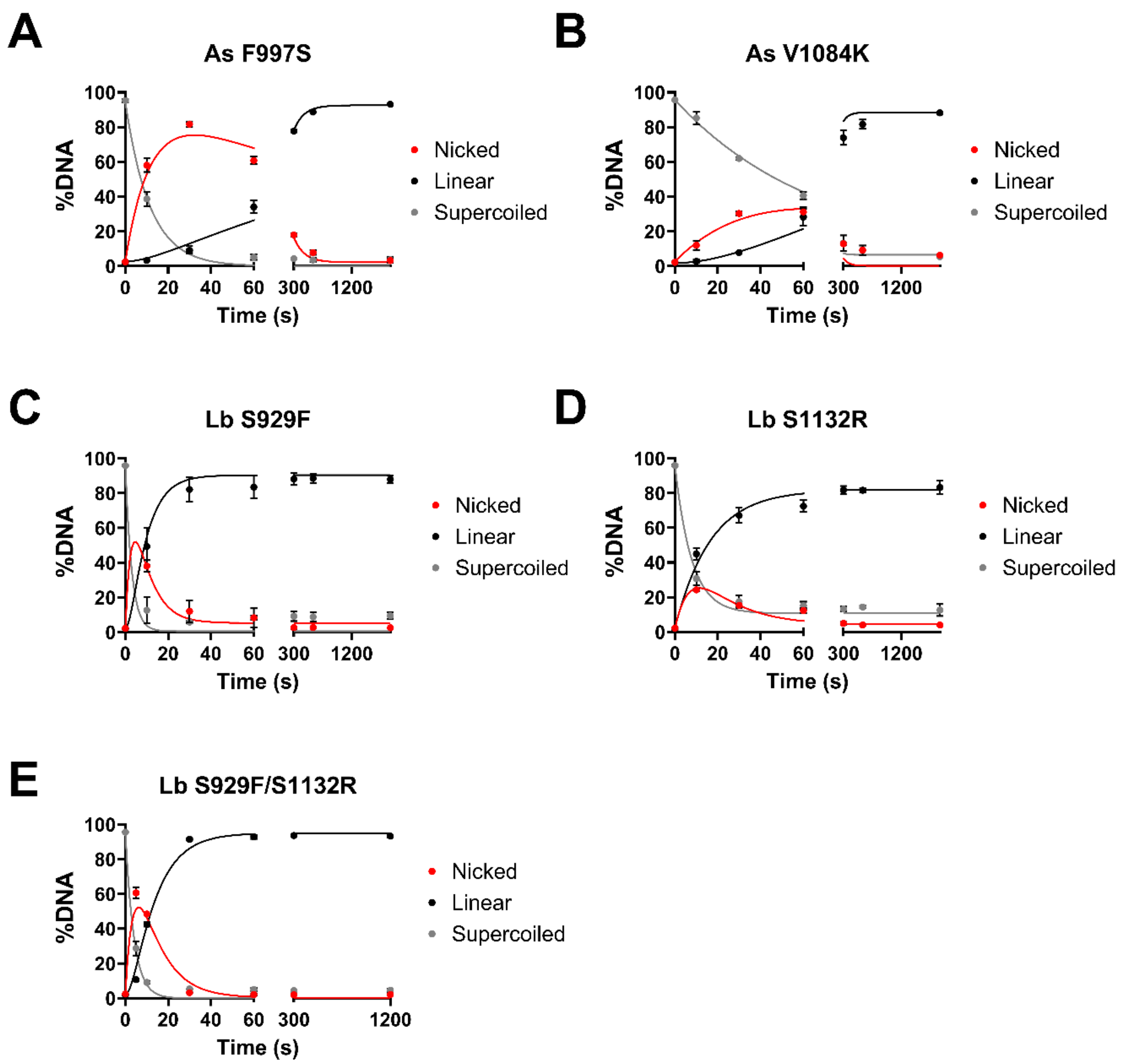


**Figure S22:** Time-course plasmid *cis* cleavage by (**A**) AsF997S, (**B**) AsV1084K, (**C**) LbS929F, (**D**) LbS1132R, (**E**) LbS929F/S1132R. Dots show mean, error bars show s.d. Lines show fit of sequential NTS and TS cleavage model. Conducted in reaction buffers with a final concentration of 50 mM NaCl.

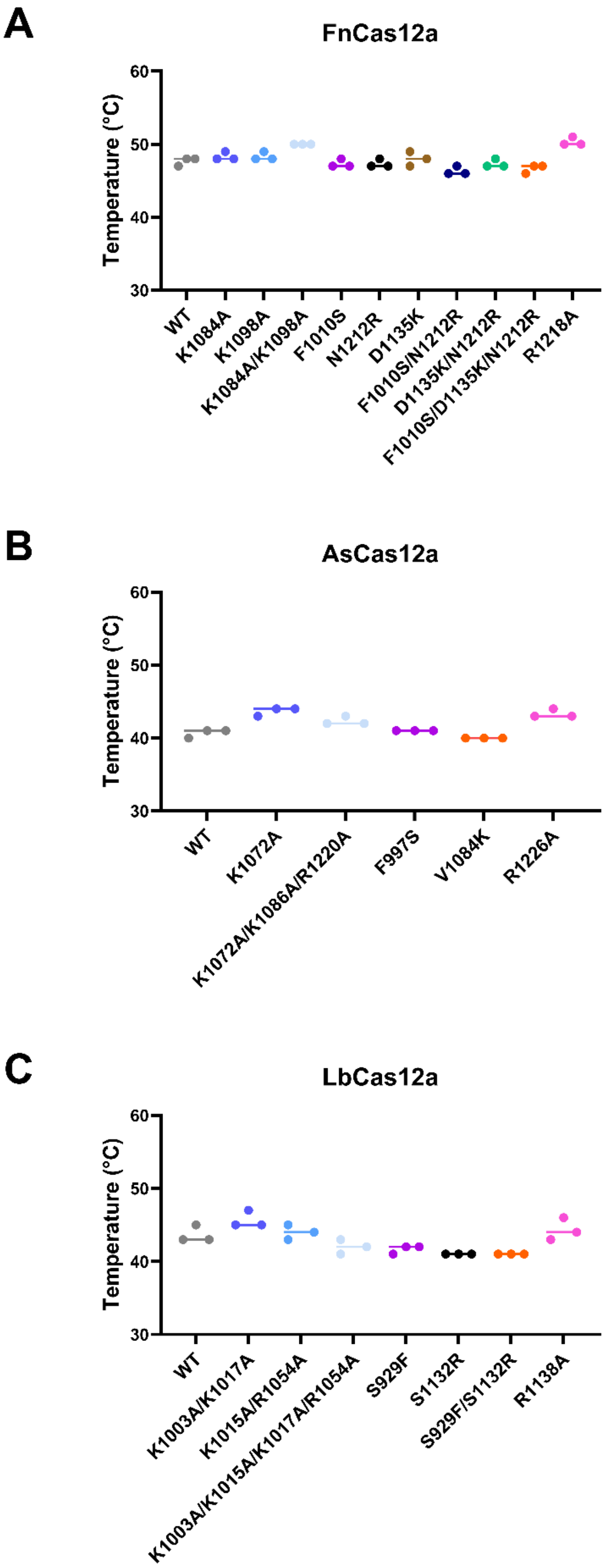


**Figure S23:** Protein thermostability, where melting temperature (°C) is defined as the fluorescence peak. Line shows mean of three replicates.

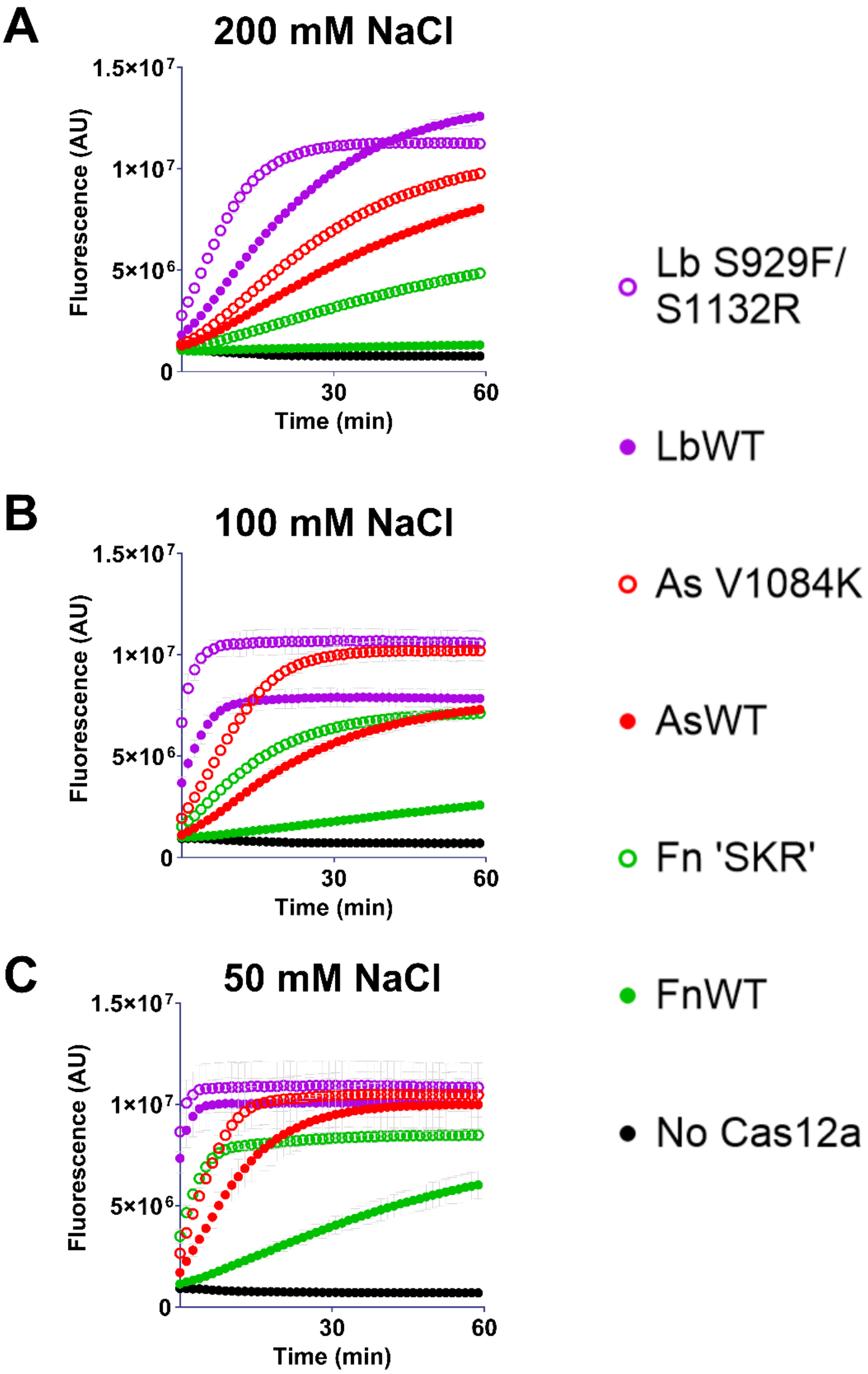


**Figure S24:** *Trans* cleavage curves of Cas12a WT and mutants (as indicated), when activated by 1 nM HPV-16 dsDNA, in reaction buffers containing a final NaCl concentration of (**A**) 200 mM, (**B**) 100 mM, and (**C**) 50 mM.

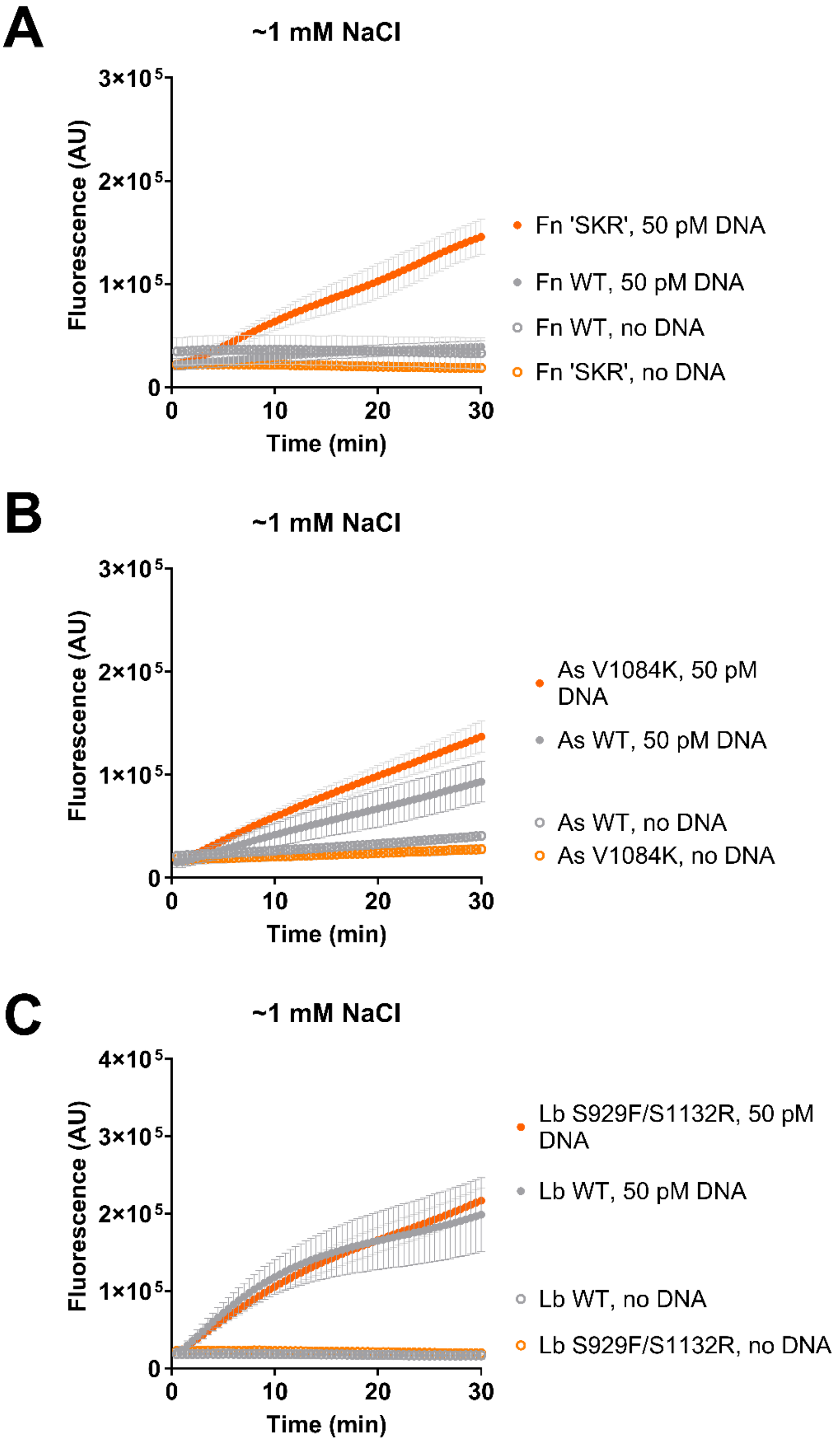


**Figure S25:** *Trans* cleavage curves in ~1 mM NaCl reaction buffer; for **A**) FnCas12a WT and 'SKR' mutant, (**B**) AsCas12a WT and V1084K mutant, and (**C**) LbCas12a WT and S929F/S1132R mutant, in the presence of either no DNA or 50 pM HPV-16 dsDNA (as indicated).

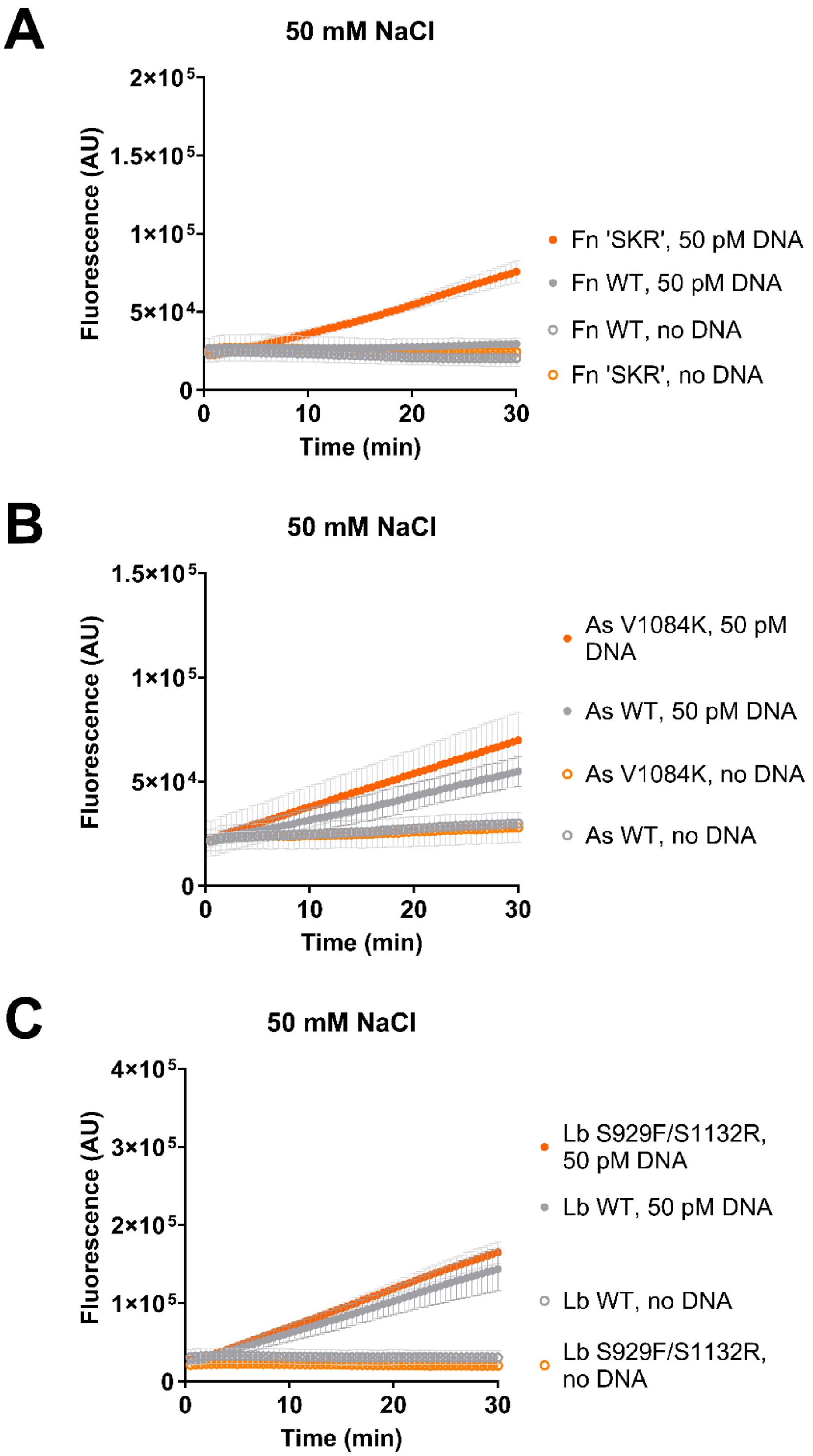


**Figure S26:** *Trans* cleavage curves in 50 mM NaCl reaction buffer; for **A**) FnCas12a WT and 'SKR' mutant, (**B**) AsCas12a WT and V1084K mutant, and (**C**) LbCas12a WT and S929F/S1132R mutant, in the presence of either no DNA or 50 pM HPV-16 dsDNA (as indicated).

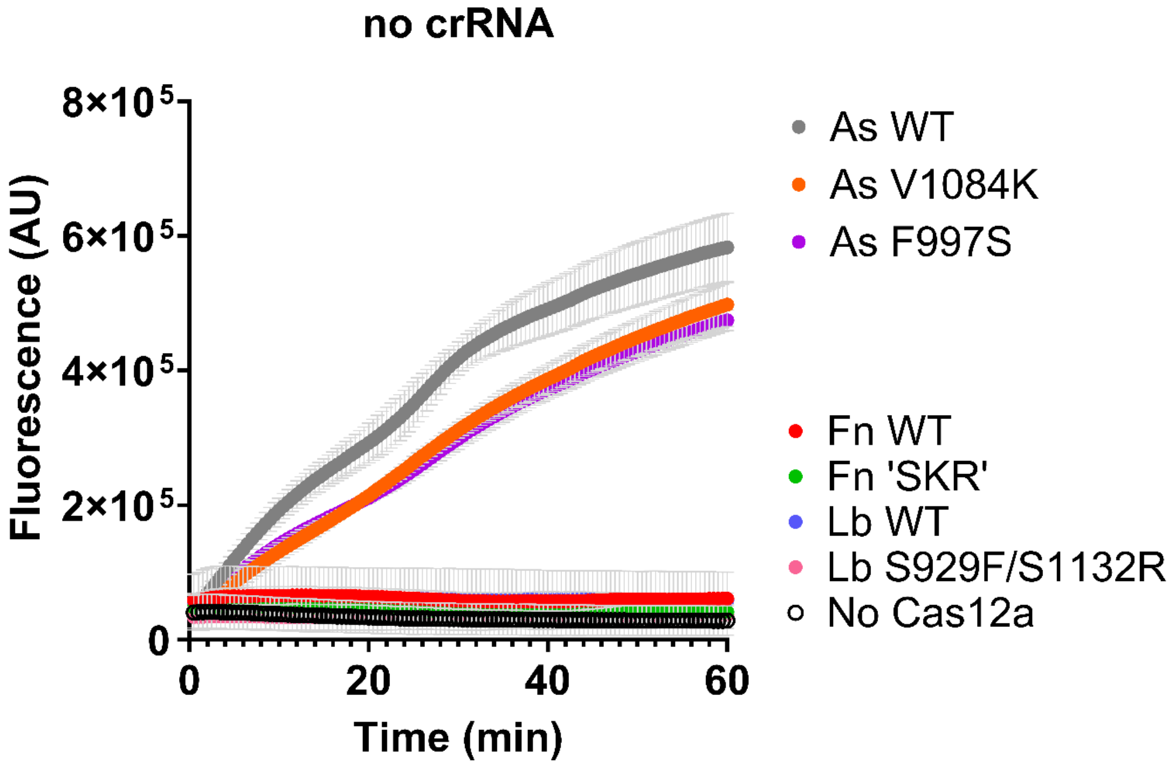


**Figure S27:** *Trans* cleavage curves of Cas12a WT and mutants (as indicated), in the absence of crRNA or any target dsDNA, reaction buffers containing a final NaCl concentration of ~1 mM.

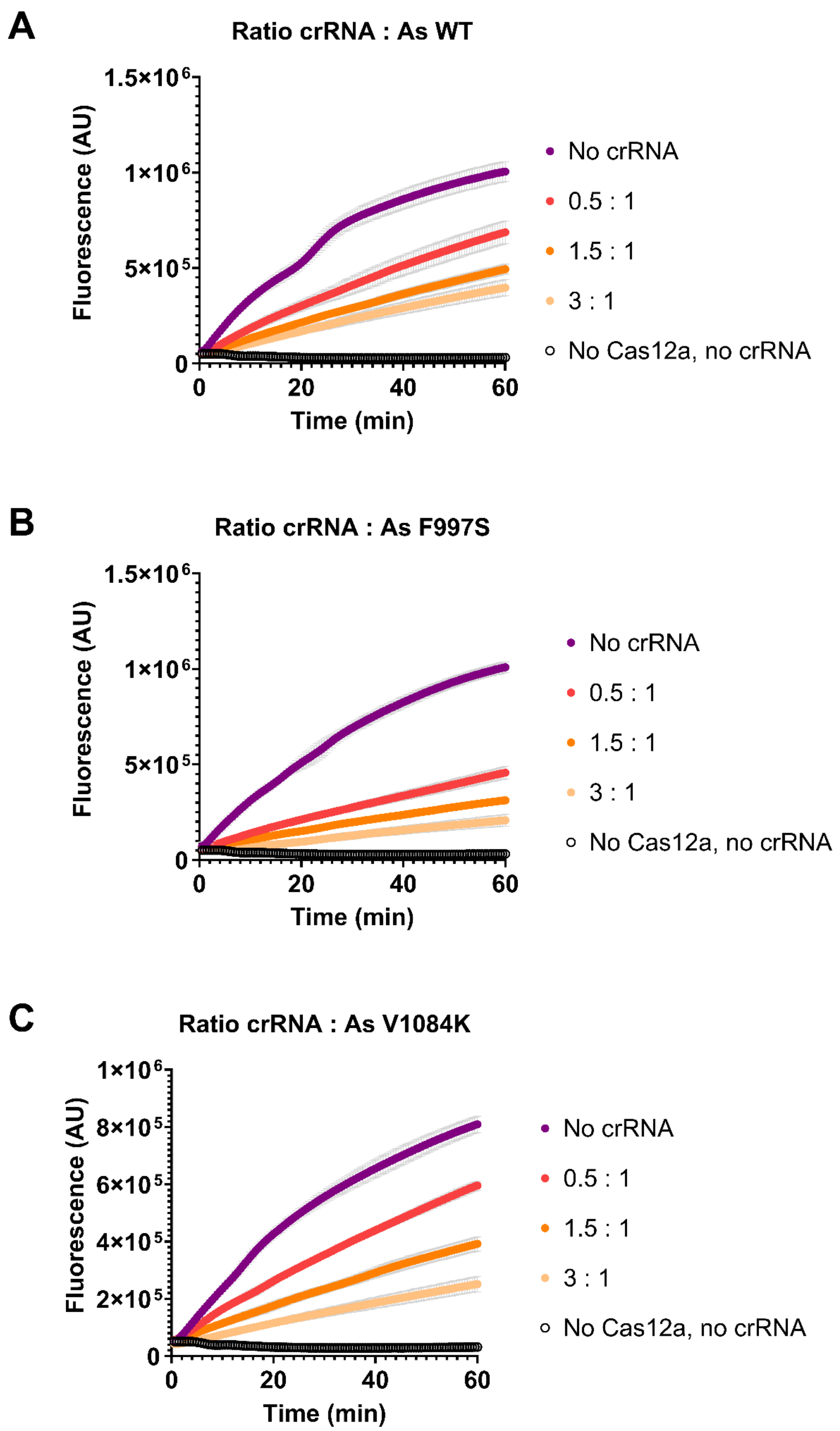


**Figure S28:** *Trans* cleavage curves of AsCas12a (**A**) WT, (**B**) F997S, and (**C**) V1084K, in the absence of any target dsDNA, and molar ratio of crRNA to Cas12a as indicated.

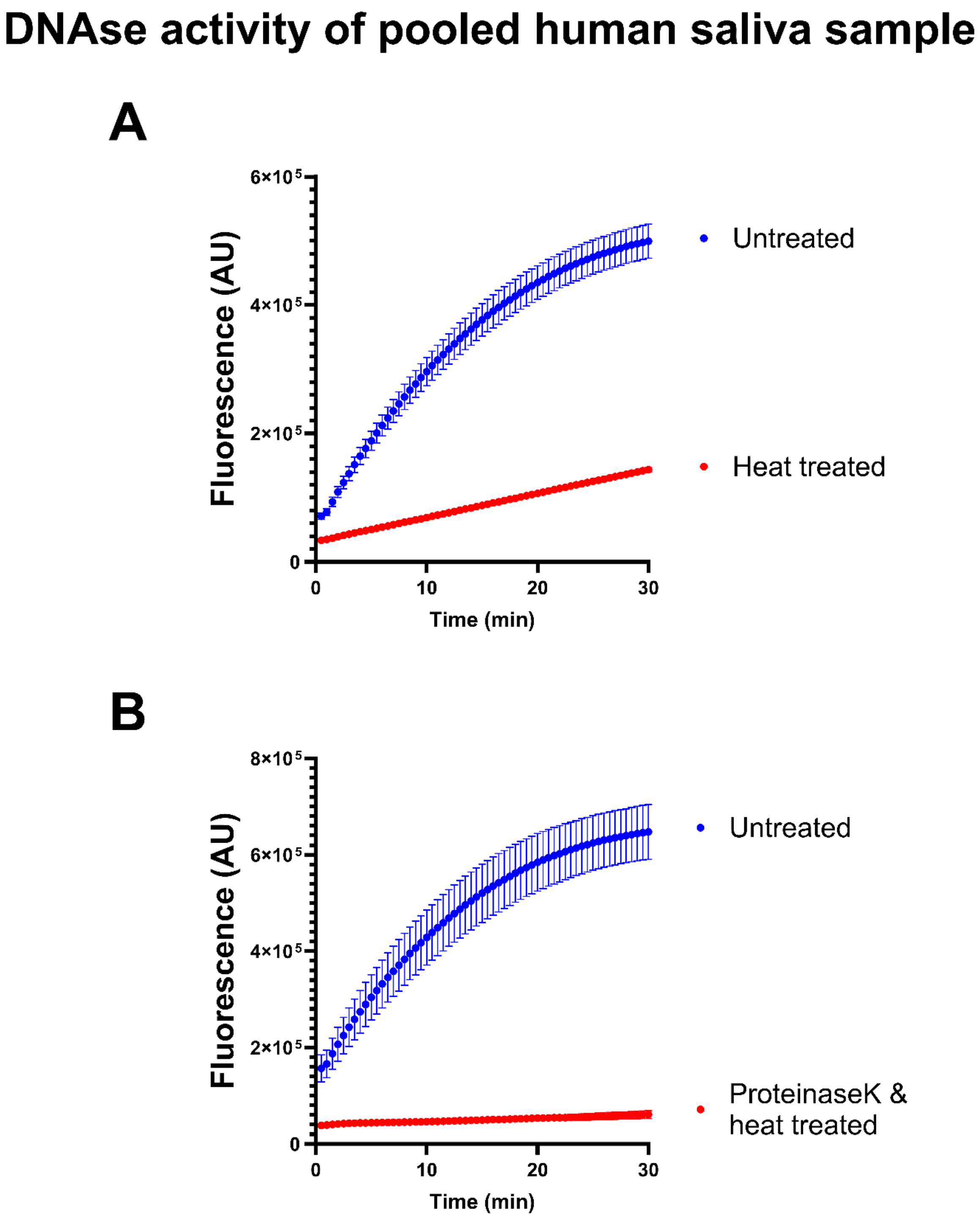


**Figure S29:** Troubleshooting the DNAse activity of pooled human saliva sample, comparing (**A**) untreated vs heat-treated at 98 °C for 10 mins, and (**B**) untreated vs addition of 2% v/v proteinase K and incubation at 55 °C for 10 mins, followed by heat-treatment at 98 °C for 10 mins.

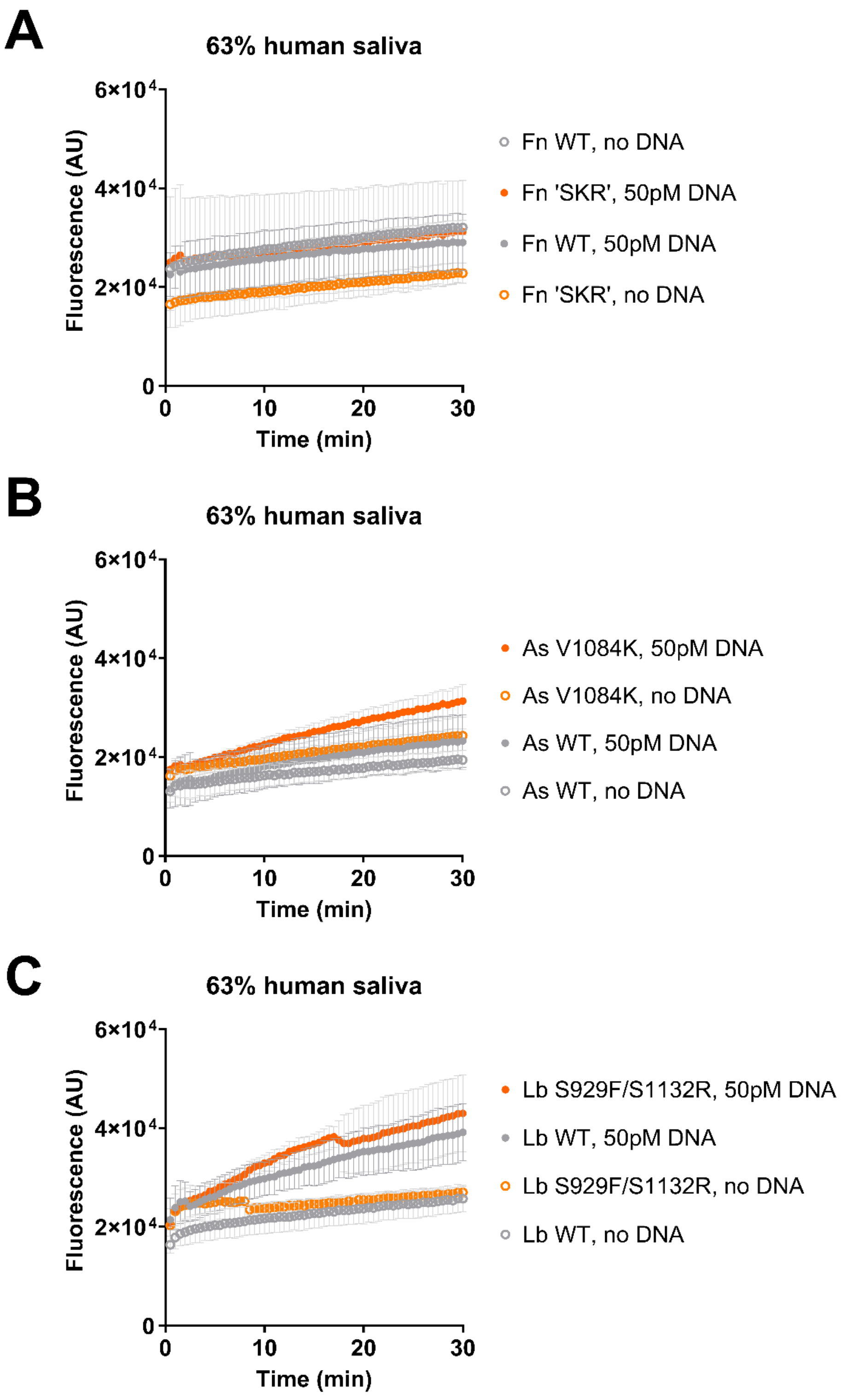


**Figure S30:** *Trans* cleavage curves in 63% human saliva sample; for **A**) FnCas12a WT and 'SKR' mutant, (**B**) AsCas12a WT and V1084K mutant, and (**C**) LbCas12a WT and S929F/S1132R mutant, spiked with either nuclease-free water or 50 pM HPV-16 dsDNA (as indicated).

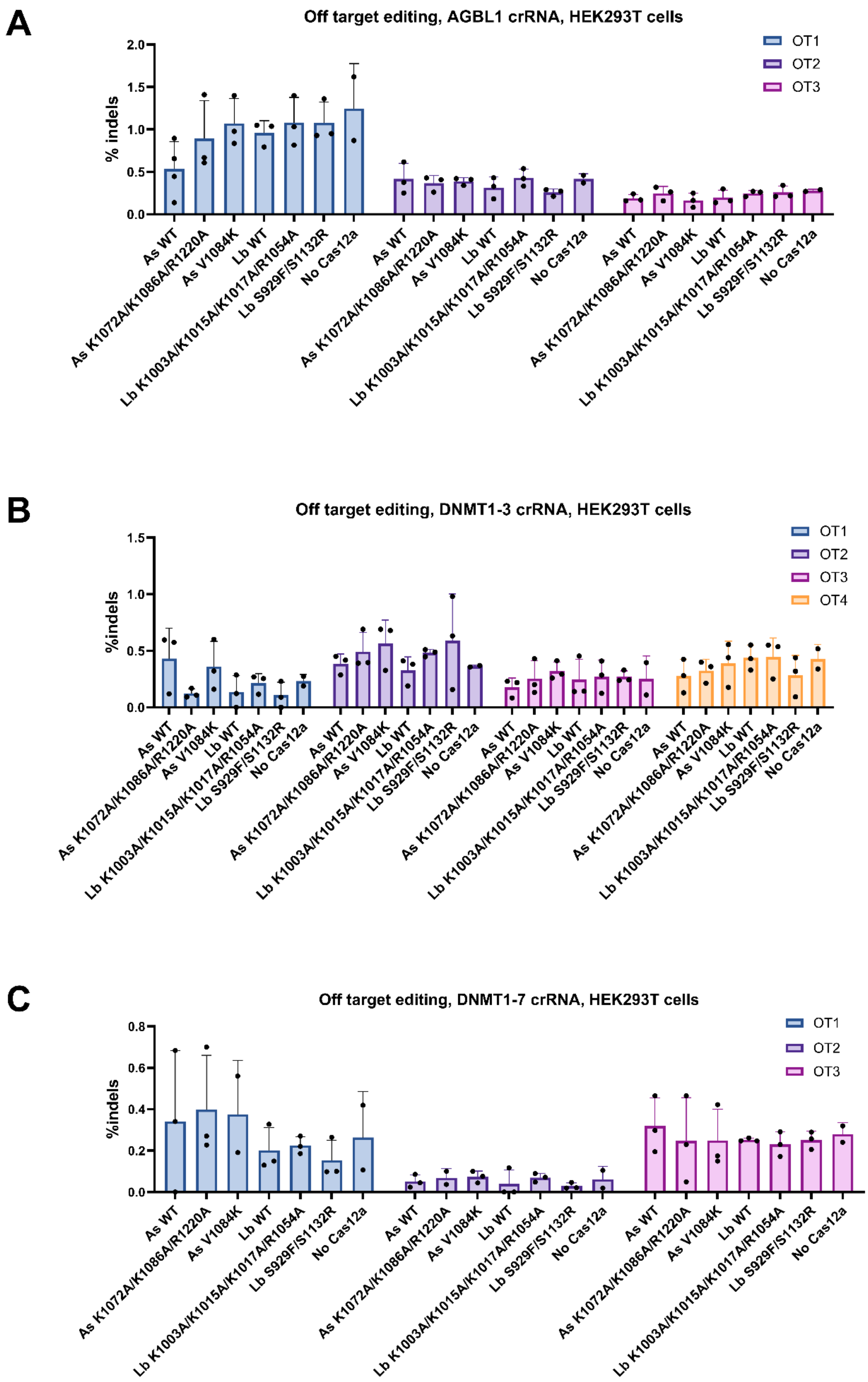


**Figure S31:** Off-target editing in HEK293T cells. Highest represented off-target (OT) sites for (**A**) AGBL1, (**B**) DNMT1-3, and (**C**) DNMT1-7, as identified in Kleinstiver 2019 (https://doi.org/10.1038/s41587-018-0011-0). Bars shown mean, error bars s.d. Statistical significance evaluated by two-way ANOVA with Tukey's multiple comparison test, p-values shown for $p<0.05$.

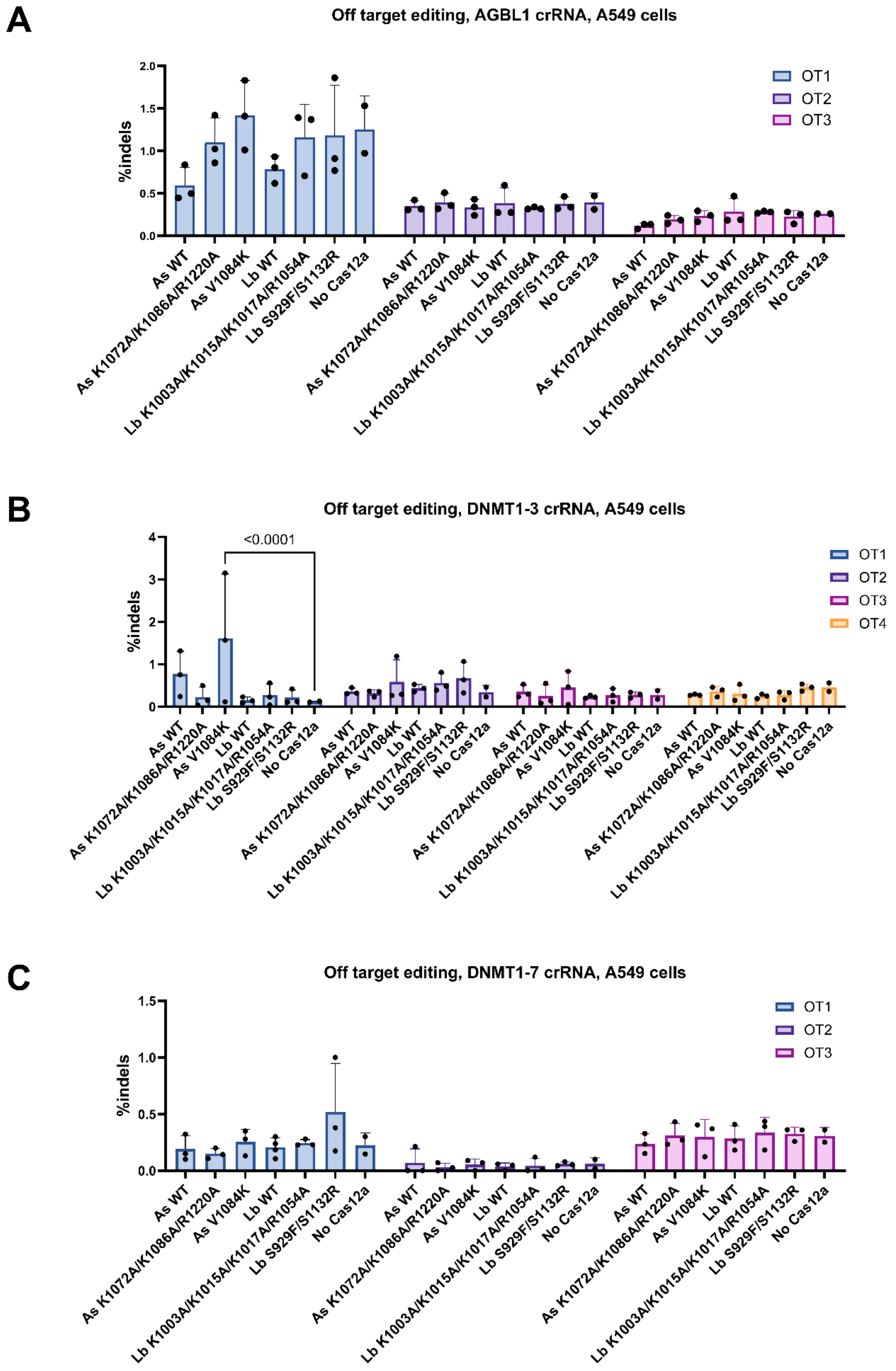


**Figure S32:** Off-target editing in A549 cells. Highest represented off-target (OT) sites for (**A**) AGBL1, (**B**) DNMT1-3, and (**C**) DNMT1-7, as identified in Kleinstiver 2019 (https://doi.org/10.1038/s41587-018-0011-0). Bars shown mean, error bars s.d. Statistical significance evaluated by two-way ANOVA with Tukey's multiple comparison test, p-values shown for p<0.05.

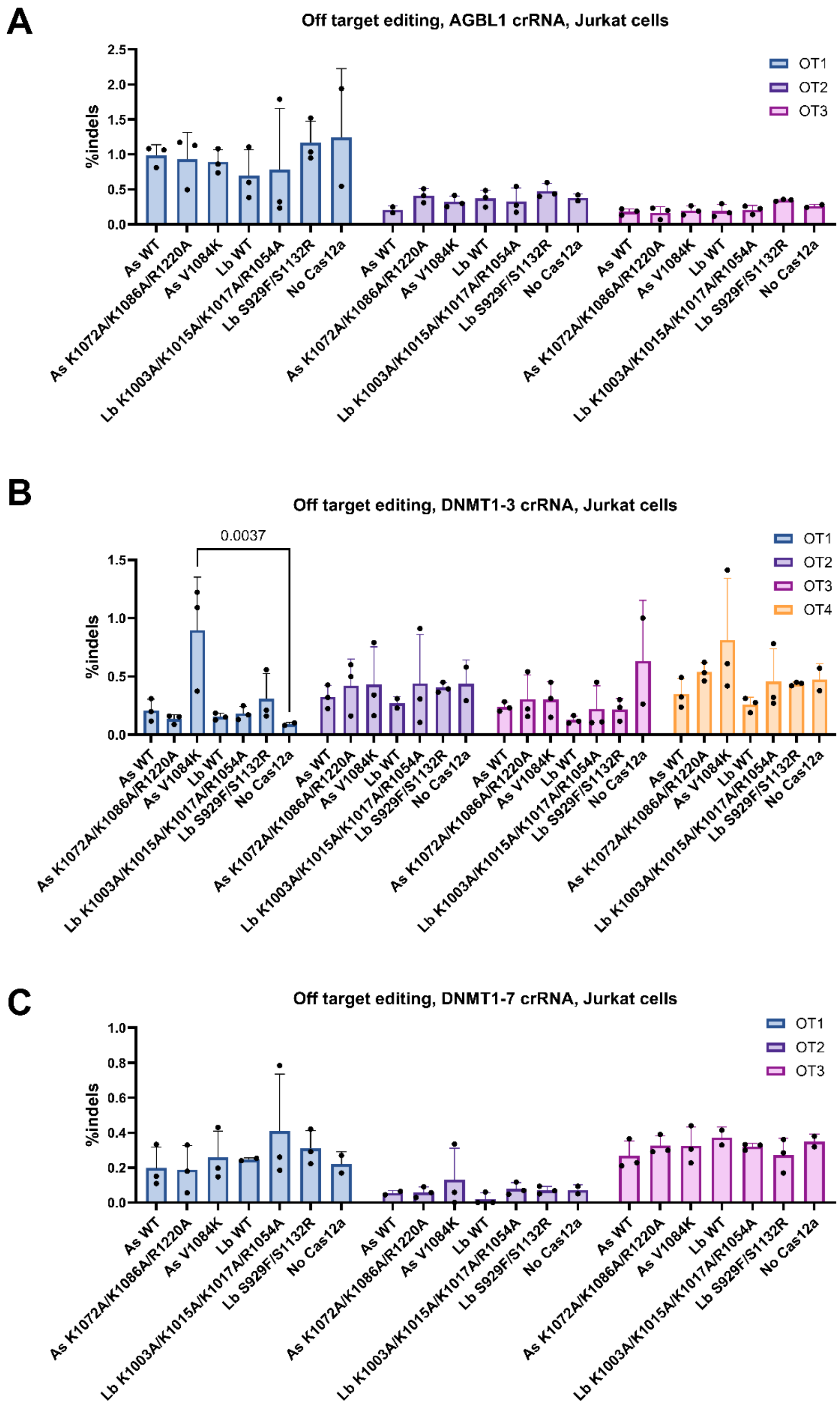


**Figure S33:** Off-target editing in Jurkat cells. Highest represented off-target (OT) sites for (**A**) AGBL1, (**B**) DNMT1-3, and (**C**) DNMT1-7, as identified in Kleinstiver 2019 (https://doi.org/10.1038/s41587-018-0011-0). Bars shown mean, error bars s.d. Statistical significance evaluated by two-way ANOVA with Tukey's multiple comparison test, p-values shown for $p<0.05$.

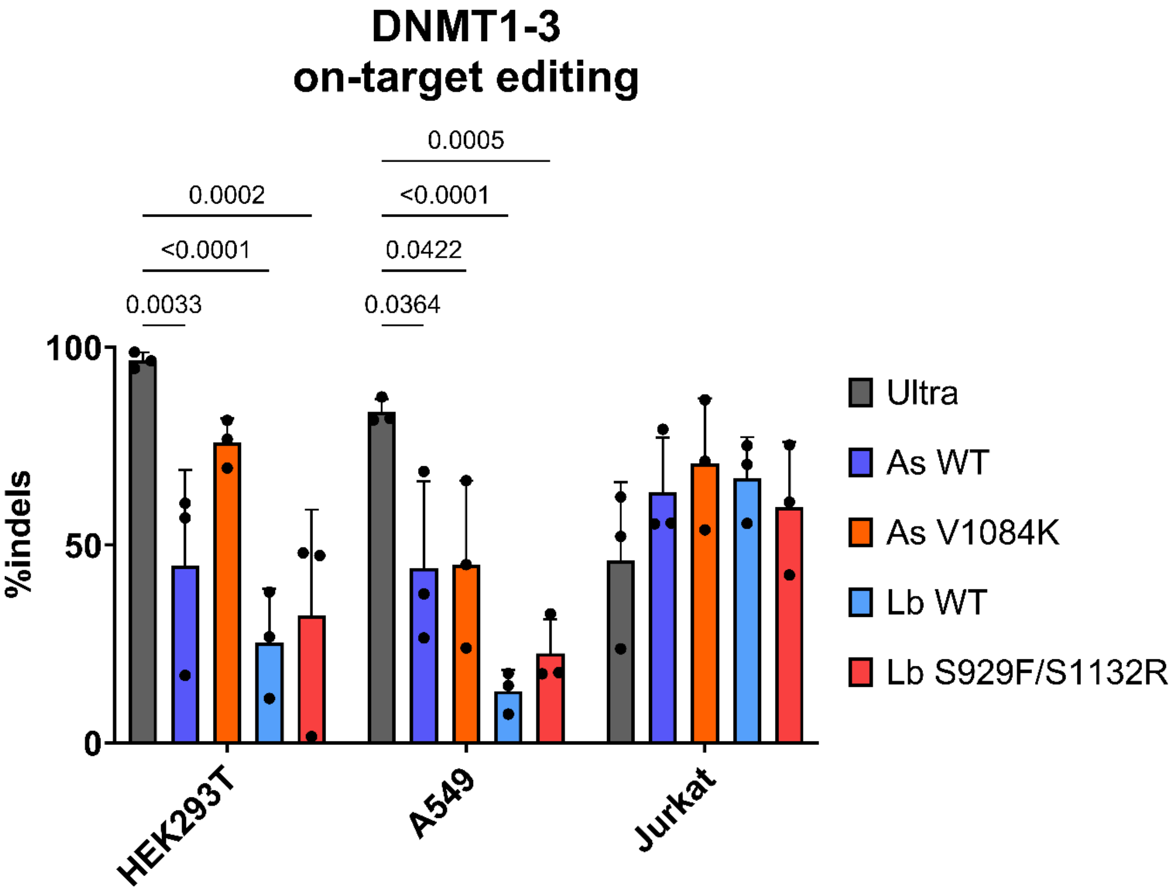


**Figure S34:** On-target editing with the DNMT1-3 crRNA, comparing 'Ultra' AsCas12a (IDT) vs nucleases made in this study, in the cell lines indicated on the x-axis. Bars shown mean, error bars s.d. Statistical significance evaluated by two-way ANOVA with Tukey's multiple comparison test, p-values shown for p<0.05.

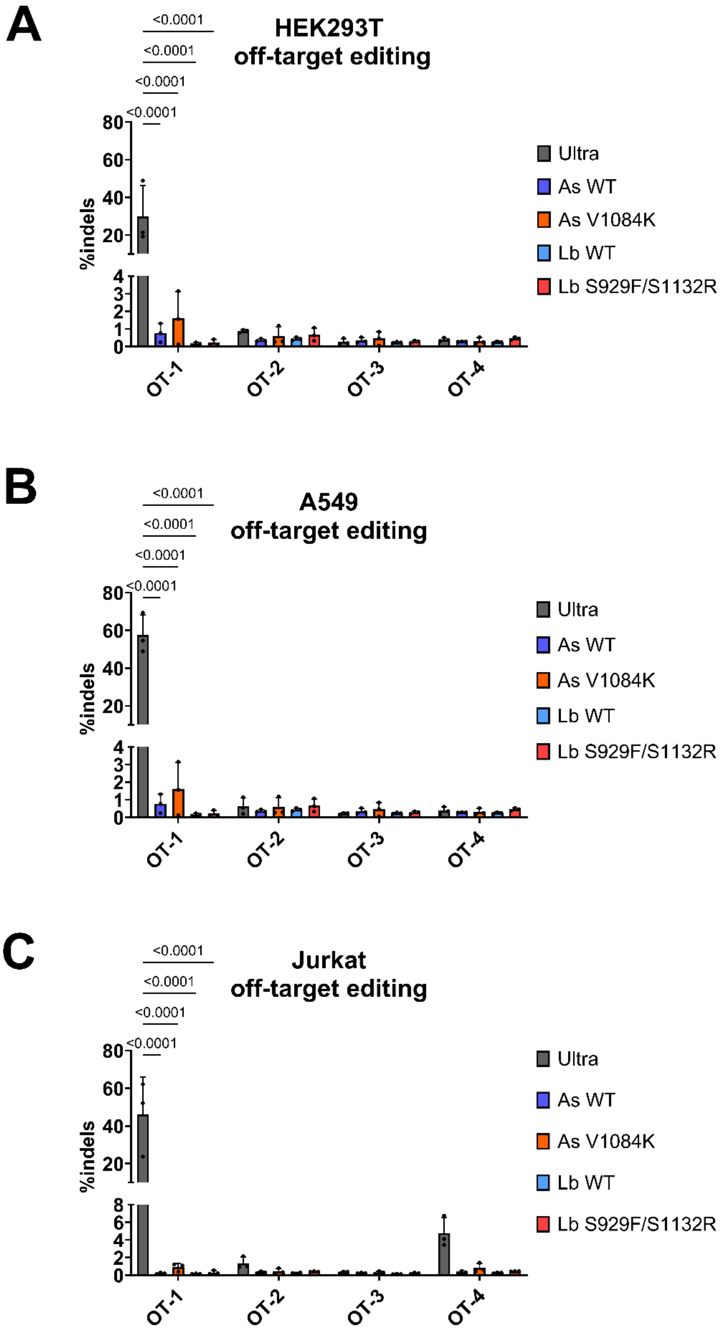


**Figure S35:** Off-target editing for DNMT1-3 crRNA, comparing 'Ultra' AsCas12a (IDT) vs nucleases made in this study, in (**A**) HEK293T, (**B**) A549, and (**C**) Jurkat cell lines. Bars shown mean, error bars s.d. Statistical significance evaluated by two-way ANOVA with Tukey's multiple comparison test, p-values shown for $p<0.05$.

| Primer name | Sequence (5' to 3') | T°(a) |
|---|---|---|
| fn_f1010s_f | gatctgaactccggctttaagag | 61 |
| fn_f1010s_r | ctcgaacaccacaatggc | 61 |
| fn_k1084a_f | attcacttccgcgatctgccccgtg | 62 |
| fn_k1084a_r | ccggctggcacatagtag | 62 |
| fn_k1098a_f | gctgtaccctgcgtatgagtcagtgagcaag | 63 |
| fn_k1098a_r | tggttgacaaagccggtc | 63 |
| fn_d1135k_f | gaacttcggcaaaaaggccgcta | 61 |
| fn_d1135k_r | ttgtaatcgaaggaaaactcg | 61 |
| fn_n1212r_f | ctcagtgctgcgtacaatcctgcag | 59 |
| fn_n1212r_r | gtcagcttggcgaaaaatttc | 59 |
| fn_r1218a_f | cctgcagatggcgaactcaaagac | 57 |
| fn_r1218a_r | attgtattcagcactgag | 57 |
| as_f997s_f | aacctgaattccggctttaagagcaagag | 68 |
| as_f997s_r | ctccagcaccaccacggc | 68 |
| as_k1072a_f | atatacatctgcgatcgatcccctgaccg | 64 |
| as_k1072a_r | ggggcaggcacgtaaaac | 64 |
| as_k1086a_f | cttcgtgtgggcgaccatcaagaatcacgagag | 67 |
| as_k1086a_r | gggtccacgaagccggtc | 67 |
| as_v1084k_f | ggacccccttcaaatggaaaaccatcaagaatcacgagagc | 69 |
| as_v1084k_r | acgaagccggtcagggga | 69 |
| as_r1220a_f | gccctgatcgcgagcgtgctgcagatgc | 66 |
| as_r1220a_r | caccatggtgtcgatggcgtgag | 66 |
| as_r1226a_f | gctgcagatggcgaactccaatgccgccac | 65 |
| as_r1226a_r | acgctgcggatcagggcc | 65 |
| lb_s929f_f | gacctgaactttggctttaagaatagccg | 69 |
| lb_s929f_r | ctccagggcgatcacggc | 69 |
| lb_k1003a_f | gctgacatccgcgatcgatccatc | 58 |
| lb_k1003a_r | caggcagggatgtaaaag | 58 |
| lb_k1017a_f | ctgctgaaaaccgcgtataccagcatcgcc | 61 |
| lb_k1017a_r | gttcacaaagccggtagatgg | 61 |
| lb_k1015a_f | gtgaacctgctggcgaccaagtatacc | 60 |
| lb_k1015a_r | aaagccggtagatggatc | 60 |
| lb_k1015a_k1017a_f | cgcgtataccagcatcgccgat | 65 |
| lb_k1015a_k1017a_r | gtcgccagcaggttcacaaagcc | 65 |
| lb_r1054a_f | gaacttctctgcgacagacgccgattac | 59 |
| lb_r1054a_r | ttatagtccagggcaaac | 59 |
| lb_r1138a_f | gctgcagatggcgaacagcatcacag | 63 |
| lb_r1138a_r | atcaggctcatcagggcc | 63 |
| lb_s1132r_f | ggccctgatgcgcctgatgctgc | 70 |
| lb_s1132r_r | ataaagctagagtagaaggccttgtcg | 70 |

**Table S1:** Primers used for Cas12a mutagenesis. T°(a) indicates annealing temperature in Q5 site-directed mutagenesis protocol.

| Oligo name | Sequence (5' to 3') |
|---|---|
| Trans cleavage reporter ssDNA | /56-FAM/tttttttt/ZEN/ttt/3IaBkFQ/ (IDT product codes) |
| TS-DNMT1 | cccggtgtcacgccacttgacaggcgagtaacagacatggaccatcaggaaacattaacgtactgatgttaacagctgacccaataagtggcagag |
| NTS-DNMT1 | ctctgccacttattgggtcagctgttaacatcagtacgttaatgtttcctgatggtccatgtctgttactcgcctgtcaagtggcgtgacaccggg |
| Precleaved-TS-DNMT1 | taacagacatggaccatcaggaaacattaacgtactgatgttaacagctgacccaataagtggcagag |
| Precleaved-NTS-DNMNT1 | ctctgccacttattgggtcagctgttaacatcagtacgttaatgtttcctgatggtccatgt |
| TS-HPV-16 | ctctgccacttattgggtcagctgttaacatcagtacgttaatgtttcctgatggtccatgtctgttactcgcctgtcaagtggcgtgacaccggg |
| NTS-HPV-16 | tccccatgtcgtaggtactccttaaagttagtatttttatatgtagtttctgaagtagatatggcagcacataatgacatatttgtactgcgtgtagtatcaacaacagtaacaaatagttggttaccccaaca |
| Lb-DNMT1-3 | uaauuucuacuaaguguagaucugaugguccaugucuguuacucg |
| As-DNMT1-3 | uaauuucuacucuuguagaucugaugguccaugucuguuacucg |
| Fn-DNMT1-3 | uaauuucuacuguuguagaucugaugguccaugucuguuacucg |
| Lb-HPV-16 | uaauuucuacuaaguguagauugaaguagauauggcagcacauaa |
| As-HPV-16 | uaauuucuacucuuguagauugaaguagauauggcagcacauaa |
| Fn-HPV-16 | uaauuucuacuguuguagauugaaguagauauggcagcacauaa |
| Lb_DNMT1-7 | uaauuucuacuaaguguagaugcucagcaggcaccugcctcagcu |
| Lb_AGBL1 | uaauuucuacuaaguguagaugauugaaggaaaaguuacaaaggu |
| As_DNMT1-7 | uaauuucuacucuuguagaugcucagcaggcaccugcctcagcu |
| As_AGBL1 | uaauuucuacucuuguagaugauugaaggaaaaguuacaaaggu |

**Table S2:** Oligonucleotide sequences used in *cis* and *trans* cleavage assays, DNA detection, and genome editing of human cell lines.

| Cas12a | $k_{NTS}$ ($s^{-1}$) | | | $k_{TS}$ ($s^{-1}$) | | |
|---|---|---|---|---|---|---|
| | **Mean** | **s.d.** | **n** | **Mean** | **s.d.** | **n** |
| Fn R1218A | 6.30e-5 | 3.47e-6 | 3 | 1.83e-4 | 1.27e-5 | 3 |
| Fn WT 200 mM NaCl | 0.284 | 0.023 | 2 | 0.254 | 0.046 | 2 |
| Fn WT 100 mM NaCl | 0.519 | 0.076 | 2 | 0.261 | 0.039 | 2 |
| Fn WT | 0.318 | 0.006 | 2 | 0.254 | 0.025 | 2 |
| Fn WT ~1 mM NaCl | 0.233 | 0.003 | 2 | 0.200 | 0.029 | 2 |
| Fn K1084A | 0.179 | 0.045 | 3 | 0.188 | 0.024 | 3 |
| Fn K1084A/K1098A | 0.188 | 0.008 | 3 | 0.093 | 0.003 | 3 |
| Fn F1010S | 0.337 | 0.200 | 3 | 0.150 | 0.016 | 3 |
| Fn D1135K | 0.229 | 0.018 | 3 | 0.302 | 0.036 | 3 |
| Fn N1212R | 0.249 | 0.117 | 3 | 0.336 | 0.078 | 3 |
| Fn F1010S/N1212R | 0.303 | 0.039 | 3 | 0.104 | 0.002 | 3 |
| Fn D1135K/N1212R | 0.131 | 0.011 | 3 | 0.240 | 0.014 | 3 |
| Fn F1010S/D1135K/ N1212R | 1.013 | 0.240 | 3 | 0.232 | 0.011 | 3 |
| As R1226A | 1.99e-4 | 6.56e-6 | 3 | 4.22e-4 | 7.80e-5 | 3 |
| As WT | 0.060 | 0.008 | 2 | 0.007 | 9.2 e-5 | 2 |
| As K1072A/K1086A/ R1220A | 0.029 | 0.008 | 3 | 0.001 | 1.70 e-4 | 3 |
| As F997S | 0.097 | 0.012 | 2 | 0.006 | 3.0e-4 | 2 |
| As V1084K | 0.013 | 0.001 | 3 | 0.011 | 0.003 | 3 |
| Lb R1138A | 2.58e-4 | 9.45e-6 | 3 | 7.85e-5 | 5.07e-6 | 3 |
| Lb WT | 0.104 | 0.025 | 3 | 0.066 | 0.021 | 3 |
| Lb K1003A/K1017A | 0.060 | 0.007 | 3 | 0.040 | 0.004 | 3 |
| Lb K1015A/R1054A | 0.070 | 0.004 | 3 | 0.057 | 0.011 | 3 |
| Lb K1003A/K1015A/ K1017A/R1054A | 0.050 | 0.015 | 3 | 0.044 | 0.010 | 3 |
| Lb S929F | 0.337 | 0.178 | 3 | 0.108 | 0.024 | 3 |
| Lb S1132R | 0.112 | 0.020 | 3 | 0.177 | 0.015 | 3 |
| Lb S929F/S1132R | 0.270 | 0.023 | 3 | 0.091 | 0.003 | 3 |

**Table S3:** Mean, s.d. and n, of kinetic values ($s^{-1}$) for NTS ($k_{NTS}$) and TS cleavage ($k_{TS}$), for the Cas12a indicated
